# Cumulative cgMLST provides increased discrimination of nested phylogenetic groups

**DOI:** 10.1101/2025.11.25.689980

**Authors:** Armen Ovsepian, Jose F. Delgado-Blas, Martin Rethoret-Pasty, Melissa J. Martin, François Lebreton, Sylvain Brisse

**Author notes:** Correspondence: Sylvain Brisse: Institut Pasteur, Biodiversity and Epidemiology of Bacterial Pathogens, 25-28 rue du Docteur Roux, F-75724, Paris, France.

## Abstract

**Background:** Core genome multilocus sequence typing (cgMLST) is a powerful method for bacterial strain genotyping. However, the size of the core genome decreases as the phylogenetic breadth of the target group increases, reducing discriminatory power. To overcome this discrimination/applicability tradeoff, here we developed a cumulative cgMLST approach, where sets of core loci conserved within nested phylogenetic entities are added. We illustrate this approach using the *Klebsiella pneumoniae* species complex (KpSC), for which a widely used cgMLST scheme (KpSC-cgMLST) comprises only 629 genes.

**Methods:** We created non-redundant cgMLST schemes for the individual species *K. pneumoniae sensu stricto* (Kpn-cgMLST scheme), and its multidrug resistant sublineages (SLs) SL147 and SL307. To extract core genes, we used 37,874 genome assemblies originating from over 80 countries worldwide. A methodology was set to filter redundant loci before importing them into the genotyping tool BIGSdb, where they were combined into schemes together with preexisting loci conserved at higher phylogenetic levels. The performance of the cumulative cgMLST schemes was evaluated on previously published datasets and on novel data from an inter-hospital outbreak of SL307.

**Results:** The Kpn-cgMLST, SL147 and SL307 schemes comprise 2752, 852, and 947 additional loci, respectively. The mean allele call rate of the novel loci was >99% in the validation datasets. Compared to the KpSC scheme used alone, pairwise allelic distances among isolates increased on average 5.6-fold using the Kpn scheme, and further by 20% and 30% using the SL147 and SL307 schemes, respectively. We demonstrate the added value of this increased discriminatory power for epidemiological analyses and show nearly equal discrimination when compared to whole-genome single nucleotide polymorphisms analysis.

**Conclusions:** The cumulative cgMLST strategy combines broad phylogenetic applicability and nearly complete genotyping resolution, expanding the utility of this harmonized approach for genomic epidemiology.

## Background

Within a bacterial species, strains can display notable diversity in terms of antimicrobial resistance, virulence, and epidemic potential (1). Comparison of strain genotypes during prospective surveillance is essential to detect common-source outbreaks, and strain genotyping is also used to match infection cases with isolates from suspected contamination sources. Therefore, accurate characterization of strain diversity has become central to surveillance and targeted public health interventions. The advent of whole genome sequencing has transformed bacterial genotyping and epidemiology, with core genome multilocus sequence typing (cgMLST) and whole-genome single nucleotide polymorphisms (SNP) analyses being the two most widely adopted sequencing-based strain genotyping approaches (2,3).

While cgMLST retains the intuitive nature of MLST (4), it offers substantially higher resolution than 7-gene classical MLST by leveraging the genetic variation of a much larger proportion of the genome (5,6). A cgMLST scheme, i.e. a standardized genotyping framework for cgMLST analysis, typically employs a few thousands of gene loci (7,8). Still, the number of loci that can be included in a cgMLST scheme depends on the targeted phylogenetic level. Schemes developed for taxonomically broad groups (e.g., entire genera or complexes of closely related species) will incorporate relatively small core genomes, comprising only the broadly shared gene loci, in order to ensure applicability across all target species. In contrast, schemes designed for phylogenetically more homogeneous groups will be able to include a larger number of gene loci, providing higher discriminatory power, while being restricted in their genotyping applicability to specific phylogroups, as these gene loci will often not be present in bacterial genomes belonging to other groups. Thus, the cgMLST approach faces a tradeoff between genotyping discriminatory power and breadth of applicability. Ideally, a genotyping method should be able to type a broad set of strains belonging to several species, while at the same time achieving high discrimination among strains for genomics-based surveillance and outbreak investigations. The need for high resolution genotyping is particularly true for the so-called high-risk clones, which are broadly distributed and sampled from multiple sources.

Here, we introduce the cumulative cgMLST strategy, which explicitly addresses the cgMLST discrimination/applicability tradeoff by integrating genotyping information from schemes designed iteratively for nested phylogenetic groups. While integrating schemes, to avoid database redundancy and multiple counts of genetic variation, gene loci that are part of phylogenetically broader schemes should be excluded from the cgMLST schemes designed for nested groups.

To illustrate the cumulative cgMLST approach, we used *Klebsiella pneumoniae* as a model. *Klebsiella* spp. is the third most common microorganism responsible for healthcare associated infections in Europe (9), and features twice (as ESBL-, and as carbapenem-producer) in the top-6 WHO priority pathogens list for which antimicrobial resistance represents a public health threat (10). *K. pneumoniae*, the medically most important species, is a leading etiological agent of neonatal sepsis worldwide (11). *K. pneumoniae* is broadly distributed ecologically, being found in a wide range of host-associated and environmental niches, and exhibits extensive phenotypic and genotypic diversity (12,13). Population analyses of *K. pneumoniae* isolates have shown that the rapid emergence of multidrug-resistant (MDR) *K. pneumoniae* is largely driven by the geographic dissemination of successful lineages or clones, including the sublineages (SLs) SL147 and SL307 (13).

While a cgMLST scheme for the *K. pneumoniae* species complex (KpSC) developed in 2014 is widely used (7,13), the relatively small number of loci (n=629) that were included in this scheme limits its discriminatory power. In this study, we aimed to complement this scheme, which forms a basis for the definition of KpSC sublineages (14), with a cgMLST scheme dedicated to *K. pneumoniae sensu stricto* (Kpn), as well as cgMLST schemes for two of its high-risk sublineages, SL147 and SL307. Sublineage SL147 includes the *K. pneumoniae* clonal group CG147 (15), which comprises extensively resistant isolates that carry multiple and diverse resistance genes and mobile genetic elements (16,17). SL307 is another prominent emerging clonal group of *K. pneumoniae,* the members of which are often multidrug-resistant and ESBL-producing (16,17). The novel cgMLST schemes are intended to be used in combination with the KpSC scheme, exemplifying the cumulative cgMLST genotyping strategy. We demonstrate the benefits of this approach for outbreak investigation.

## Methods

### Identification of candidate loci for the core genome MLST schemes

Selected genome assemblies were used to identify candidate loci for the core genome MLST schemes with chewBBACA v3.3.9 (18). The available prodigal training file for *K. pneumoniae* (19) and the following functions available within the chewBBACA suite were used: i) Whole genome multi-locus sequence typing (wgMLST) scheme creation with default settings (BLAST score ratio of 60%, coding sequence size variation threshold of 20%); ii) Allele calling using the wgMLST scheme; iii) Definition of the cgMLST scheme (95% threshold). Paralogous loci identified during allele calling (paralogous_loci.tsv output file) were removed; iv) Allele call and schema evaluation. From the resulted cgMLST loci with a 95% threshold, loci having high frequency (≥1%) of non-informative paralogous hits (assigned as NIPH or NIPHEM during allele calling) were removed. The remaining loci were compared to the loci of the existing cgMLST scheme for *Klebsiella pneumoniae* species complex (n=629) (7,14) and to the *Klebsiella pneumoniae* rMLST loci (n=53) (20). In the case of the sublineages SL147 and SL307, the new Kpn-cgMLST loci were additionally included in the comparison. All the alleles of the new loci were compared with allele 1 of the existing cgMLST scheme loci and the alleles of the rMLST reference *Klebsiella pneumoniae* DSM 30104 (assembly GCA_000281755.1) for the rMLST loci, with BLASTn with default parameters. With thresholds set at ≥80% for identity and ≥80% for query or subject coverage, matching loci were removed. The remaining loci were integrated into the BIGSdb-Pasteur database to test their allele call rate using this tool.

### Implementation in the BIGSdb-Pasteur KpSC database

To define a type allele for each locus, the allele with the highest frequency among the setup isolates was used. Then, loci were created in the BIGSdb-Pasteur KpSC database with a standard length defined as the type allele length, while maximum and minimum accepted allele lengths were defined as the type allele length ±5%, rounded to the nearest integer that is multiple of 3. The allele call rates were first assessed using the same assemblies which were initially used for the identification of the candidate loci with chewBBACA. Subsets of the total alleles that encompass the known diversity within a locus, were defined as exemplar alleles (find_exemplars.pl BIGSdb script) to speed up the locus scanning process (see BIGSdb documentation). The assemblies were searched for existing and novel alleles in three steps: (i) first tagged for the existing alleles (autotag.pl script with the following parameters: --word_size 30, --fast, --exemplar), then scanned for new alleles (scannew.pl script with the following parameters: --word_size 30, --fast, --coding_sequences, --alignment 90 and --identity 90 --type_alleles, --exemplar), and finally re-tagged with the same parameters to attribute novel alleles to the genomes. The type_alleles argument in the scannew.pl script ensured that only alleles with ≥90 percent identity and ≥90 percent alignment compared to the type alleles are accepted. To retain only loci that are present in most genomes, the loci that were tagged in ≤90% of the tested isolates were excluded from the final scheme. The final number of loci that were included in the new Kpn-, SL147-, and SL307-cgMLST schemes were 2,752, 852, and 947, respectively.

To name the new Kpn-cgMLST loci created in the database, the type allele sequences of each locus were aligned against the genome of *K. pneumoniae* NTUH-K2044 assembly ASM988v1 by using blastn with default options. The loci that were present on the reference genome with ≥80% identity and coverage were named based on the order of their occurrence (5’ end) on the reference genome, incrementally (e.g., Kpn0001, Kpn0002 up to Kpn2743). Loci not identified on the reference genome were assigned arbitrary names beginning from Kpn2744 onwards. For both the SL147 and SL307 schemes, all loci were sequentially assigned arbitrary identifiers starting from SL001. For the typing of the bacterial isolates in the validation datasets, the same autotag.pl and scannew script parameters were used as those used during the testing of the scheme.

Finally, for Kpn-cgMLST loci we checked whether allele calls were robust to sequencing coverage depth. We searched for non-reproducible allele calls at low assembly coverage depths, by randomly selecting read pairs simulating various coverage depths (10X, 20X, 30X, 40X, 50X, 60×, 70×, 80×, 90×), as drawn from the fastq files of the isolates SB5806, SB5734 and SB5514, and assembled them with the fq2dna pipeline (21), in triplicate for each coverage. For all the tested genomes, the numbers of missing allele calls were similar at 90×, 80×, 70×, 60×, and 50× coverage, while at coverage levels below 50×, the number of missing allele calls increased slightly (**Supplementary Figure S1**). From all the loci that were not missing in ≥ 2 samples with 90×, 80× and 70× coverage (*i.e*., were conserved), no locus was found to be consistently missing in ≥ 2 samples with 40× and 50× coverage in the tested isolates. Therefore, allele calling for all the 2,752 loci was considered reproducible down to 40× coverage.

### Gene and gene product annotations

Gene and gene product annotations were assigned to the novel core genome loci with the bioinformatics tool bakta (22). To this aim, the prodigal training file for *Klebsiella pneumoniae* utilized above for cgMLST scheme development was used, and as genus and species parameters the terms “Klebsiella” and “pneumoniae” were used for the annotation of gram-negative bacteria. The annotations were included in the fields “Common name” and “Product” of the locus definitions in the database.

### Bacterial genomes included in the study

The following datasets were used for this study:

#### Dataset 1

A dataset for the development of the core genome MLST scheme for *K. pneumoniae sensu stricto* (Kpn-cgMLST). Initially, 37,874 publicly available *K. pneumoniae* assemblies, which originate from isolates that belong to the phylogroup Kp1 (14), were downloaded from the BIGSdb-Pasteur KpSC database (5) on the 11^th^ of September 2024. The quality of the assemblies was checked with QUAST v5.0.2 (23). Only genomes fulfilling the following criteria were considered for further analysis: Total number of contigs ≤ 198 (corresponding to the mean plus twice the standard deviation), genome total length within the range of 5,565,664 bp ± 184,244 bp (mean ± twice the standard deviation), GC% within the range of 57.15 % ± 0.24 % (mean ±twice the standard deviation), assembly N50 value ≥ 60 kbp, assembly L50 value ≤ 30, number of ambiguous nucleotides ≤ 30 per 100 kbp. These values are consistent with the klebNET-GSP genome assembly quality criteria (24). Additionally, 19 *K. rhinoscleromatis* and *K. ozaenae* isolates, which have the KpSC cgMLST-based LIN code prefixes 0_0_442, 0_0_443 and 0_0_439 (14), were also removed. From the 32,234 assemblies that passed the quality control, 2,483 assemblies covering the whole range of unique LIN code size 4 prefixes (i.e. the first four of the ten LIN code levels) and therefore uniquely representing all clonal groups (14), were chosen. In cases when the LIN code size 4 prefixes were present in more than one isolate, the isolate with the lowest id in the BIGSdb-Pasteur database were chosen. These 2,483 isolates come from 82 different countries across Europe (n=1,113), Asia (n=595), North America (n=316), Africa (n=242), Oceania (n=103), and South America (n=73). These assemblies were used to identify candidate loci for the Kpn-cgMLST scheme. Check section “Availability of data and materials” for a link to retrieve the data.

#### Dataset 2

We next created a dataset for the development of the cgMLST scheme for *K. pneumoniae* sublineage SL147. From the 32,234 publicly available BIGSdb-Pasteur KpSC assemblies that passed the quality control, 1,069 *K. pneumoniae* SL147 assemblies (defined with LIN code prefix 0_0_197) covering the whole range of unique full LIN-codes were chosen. Check section “Availability of data and materials” for a link to retrieve the data.

#### Dataset 3

We next created a dataset for the development of the core genome MLST scheme for *K. pneumoniae* sublineage SL307. From the 32,234 assemblies that passed the quality control, 1,080 *K. pneumoniae* SL307 assemblies (defined with LIN code prefix 0_0_369) covering the whole range of unique full LIN-codes were chosen. Check section “Availability of data and materials” for a link to retrieve the data.

#### Dataset 4

This dataset was created to test the performance of the Kpn-cgMLST scheme at low coverage depths. This dataset included three genome sequences found in BIGSdb-Pasteur with ids 6302, 6232, 4560 corresponding to the isolates SB5806, SB5734, SB5514, respectively.

#### Dataset 5

This dataset was used as validation dataset for the Kpn-cgMLST scheme, using 1,969 *K. pneumoniae* isolates from 455 hospitals in 36 European countries (EuSCAPE project) described in Grundmann *et al.* (25).

#### Dataset 6

This dataset was used as validation dataset for the Kpn-cgMLST scheme, with 1,705 *K. pneumoniae* isolates from clinical, community, animal and environmental settings from Italy (SpARK project) described in Thorpe *et al.* (26).

#### Dataset 7

This dataset was used as validation dataset for the Kpn-cgMLST scheme, with 2,520 *K. pneumoniae* Norwegian isolates from human, animal and marine sources described in Hetland *et al.* (27).

#### Dataset 8

This dataset was used as validation dataset for the Kpn-cgMLST scheme, with 1,587 *K. pneumoniae* isolates from 16 different countries across Asia (n=945), South America (n=305), Africa (n=195), North America (n=98), and Europe (n=19), collected from 2003 until 2021 by the Multidrug-Resistant Organism Repository and Surveillance Network (**Supplementary Table S1**) (PRJNA1354878).

#### Dataset 9

This dataset was used as validation dataset for the SL147-cgMLST scheme, comprising all the 1,404 public BIGSdb-Pasteur KpSC database SL147 isolates, as of 1^st^ of July, 2025.

#### Dataset 10

This dataset was used as validation dataset for the SL307-cgMLST scheme, with all the 976 public BIGSdb-Pasteur *K. pneumoniae* SL307 isolates, as of 1^st^ of July, 2025.

#### Dataset 11

This dataset was used as validation dataset for the SL147-cgMLST scheme, as the ST147 outbreak dataset from Italy, with 114 isolates described in Martin *et al.* (28).

#### Dataset 12

Finally, this dataset was used as validation dataset for the SL307-cgMLST scheme, representing a ST307 outbreak dataset from a single U.S. hospital with isolates collected from 2019 until 2025, including 13 human and 35 environmental isolates (PRJNA1355201).

### Isolate collection and whole-genome sequencing

For the isolates belonging to datasets 8 and 12, DNA was extracted using the DNeasy UltraClean 96 Microbial Kit (Qiagen, Germantown, MD, USA) and libraries were constructed using the KAPA Hyperplus Library preparation kit (Roche Diagnostics Indianapolis, Indiana, USA). Libraries were quantified using the KAPA Library Quantification Kit – Illumina/Bio-Rad iCycler™ (Roche Diagnostics) on a CFX96 real-time cycler (Biorad, Hercules, CA, USA). Libraries were normalized to 2nM, pooled, denatured, and diluted to 1nM. Whole genome sequencing was performed using a MiSeq, NextSeq 500 or NextSeq 2000 Benchtop Sequencer (Illumina Inc., CA, USA) with MiSeq Reagent Kit v3 (600 cycle; 2 x 300bp), NextSeq Reagent Kit 500/550 v2 (300 cycle; 2 x 150bp), or NextSeq 1000/2000 P2 Reagents (300 Cycles) v3 kit. (Illumina, San Diego, CA, USA). bbduk v38.96 (29) was used to remove barcode and adapter sequence as well as to perform quality trimming (ktrim=“r”, k=“23”, mink=“11”, hdist=“1”, qtrim=“r”, trimq=“15”, minlen=“100”). Kraken2 v2.1.2 (30) was used for initial taxonomic assignment (top hit=“1”, undetermined reads=“<10%”) and to screen for contamination (2+ genus level hits >5%=“isContaminated”). *De novo* draft genome assemblies were produced using shovill v1.1.0 (31) with coverage estimates generated using bbmap v38.96 (32). Minimum thresholds for contig size and coverage were set at 200 bp and 49.5+, respectively.

### Phylogenetic and population structure analysis

For the *K. pneumoniae* ST307 outbreak isolates (dataset 12, PRJNA1355201), a recombination-free SNP phylogeny was created. SNP calling was performed with Snippy v.4.4.5 (33) with custom parameters (--mincov = 10; --minfrac = 0.8; and –minqual = 100), error corrected [Pilon v1.23] (34) and annotated (22) draft assemblies with the earliest isolate (by date) set as a reference. The core SNP alignment (5,469,960 bp) was filtered for recombination using Gubbins v2.3.156 (35). SNP distances were obtained from the resulting alignment of 97 core polymorphic sites by using snp-dist (36) and a phylogeny was created by inferring a maximum-likelihood tree with RaxML-NG v0.9.0 (37) using the GTR+G model and 50-50 parsimony and random starting trees.

For the visualization of core genomic relationships among the bacteria based on cgMLST analysis, Grapetree (38) was employed. GrapeTree implements a minimum spanning tree algorithm to reconstruct genetic relationships in Newick format.

For the clustering of the isolates based on the SNP or cgMLST distances, single linkage clustering was used, while the adjusted rand index was used to evaluate the similarity between clusterings.

### Statistical analysis

To assess the relationship between pairwise allelic distances computed by cgMLST and the single nucleotide polymorphism (SNP) distances, we applied linear regression using the least squares method. The Pearson correlation coefficient was calculated to quantify the strength and direction of the linear relationship between these two distance measures.

## Results

### Conceptualization of the cumulative cgMLST approach

We devised an iterative cgMLST scheme creation approach, which we propose to designate as “cumulative cgMLST”. As illustrated in **Fig. 1**, the cumulative cgMLST approach is based on the development of cgMLST schemes at multiple phylogenetic levels (e.g., species complex, species, and sublineage). To avoid redundancy, a filtering step is performed, during which loci that are part of upper-level schemes are excluded from those designed for phylogenetically nested groups. After integrating the non-redundant loci into a unique database, different combinations of loci (schemes) can be applied for cgMLST genotyping, depending on the isolates under investigation. In our example on *K. pneumoniae*, we expanded the existing species complex cgMLST scheme (KpSC-cgMLST) by developing an additional scheme for *K. pneumoniae sensu stricto* (Kpn-cgMLST), as well as for two of its sublineages, SL147 (SL147-cgMLST) and SL307 (SL307-cgMLST).

**Fig. 1.**
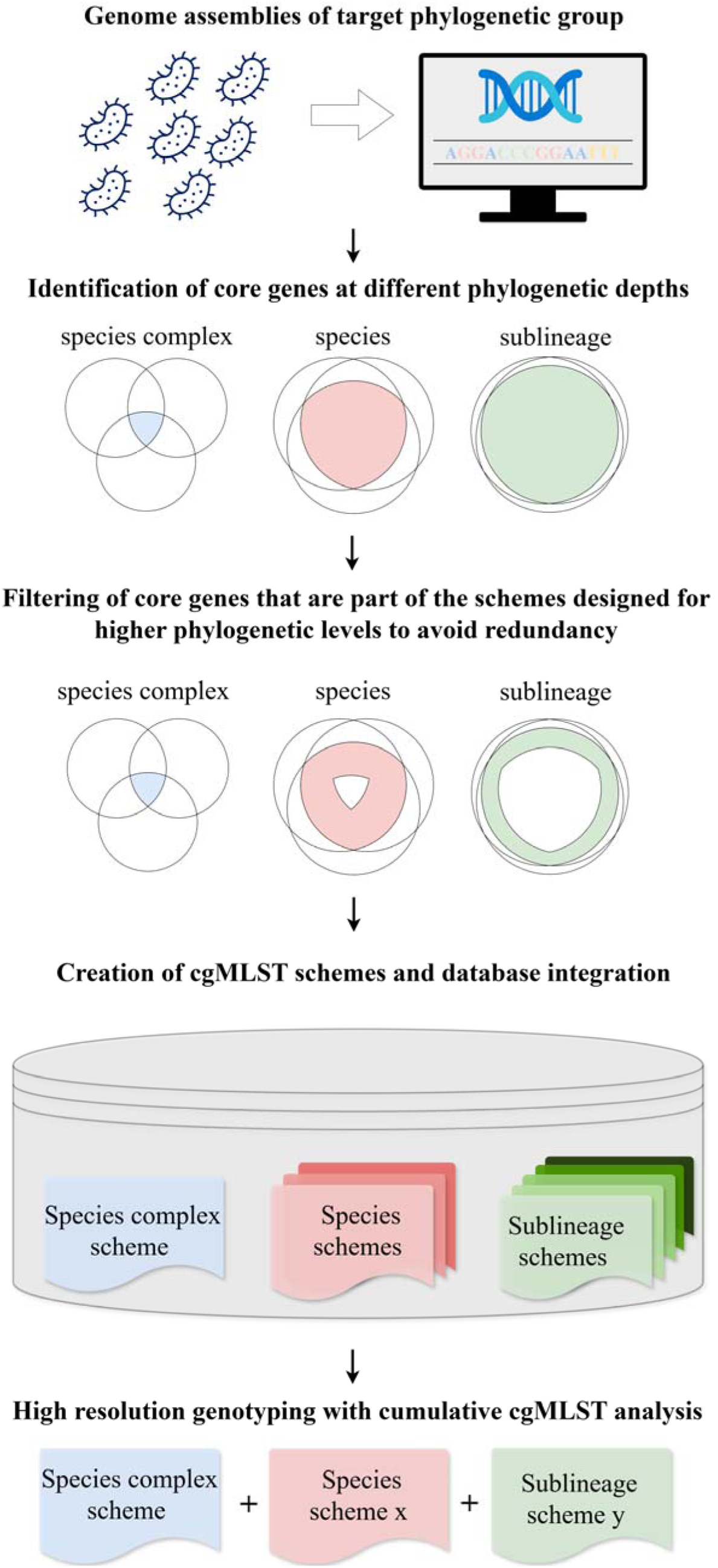
Schematic presentation of the cumulative cgMLST strategy.

### Definition and evaluation of the Kpn-cgMLST scheme

The workflow with the main steps for the development of the Kpn-cgMLST scheme can be found in **Supplementary Figure S2**. The resulting Kpn-cgMLST scheme includes 2,752 protein-coding gene loci, none of which overlaps with the loci present in the previously established and more broadly applicable KpSC-cgMLST (7,14) or rMLST (20) schemes. Of note, the Kpn-cgMLST scheme contains the full coding sequences of the loci found in traditional 7-loci MLST (*gapA*=996 nt, *infB*=2691 nt, *mdh*=939 nt, *pgi*=1650 nt, *phoE*=1053 nt, *rpoB*=4029 nt, *tonB*=747 nt), which are restricted to internal portions of these coding sequences (*gapA*=450 nt, *infB*=318 nt, *mdh*=477 nt, *pgi*=432 nt, *phoE*=420 nt, *rpoB*=501 nt, *tonB*=414 nt) (39). Therefore, the 7 MLST loci should not be used in combination with the Kpn-cgMLST loci for genotyping.

The Kpn-cgMLST scheme, together with the KpSC-cgMLST (n=629 loci) and rMLST (n=53 loci) schemes, totals 3,434 loci, collectively covering approximately 66% of the protein-coding sequences in the *K. pneumoniae* NTUH-K2044 reference genome (NCBI RefSeq assembly GCF_000009885.1; 5,174 CDS).

Based on the scheme setup dataset of 2,483 genomes (dataset 1), locus lengths varied from 90 to 4,950 bp (**Supplementary Figure S3**), while the total allele numbers ranged from 12 to 1314 per locus, illustrating the high level of genetic diversity of the core genes within *K. pneumoniae*. As expected, the number of alleles per locus increased with locus size (**Supplementary Figure S3**). The locus names, the positions on the reference genome, and other characteristics of the loci are presented in **Supplementary Table S2**.

The allele call rate of the Kpn-cgMLST scheme loci was determined for the BIGSdb allele calling functions, using the 2,483 scheme setup isolates, the 1,969 isolates from 36 European countries from the EuSCAPE project (dataset 5), the 1,705 isolates from clinical, community, animal and environmental settings from Italy from the SpARK project (dataset 6), the 2,520 Norwegian isolates from human, animal and marine sources collected from 2001 until 2020 (dataset 7), and the 1,587 isolates from 16 different countries across the world, collected from 2003 until 2021 by the Multidrug-Resistant Organism Repository and Surveillance Network (dataset 8) (**Table 1** and **Supplementary Figure S4**). The mean call rate per profile for the setup isolates was 99.2 ± 0.8 %, with only 13 isolates (0.5 %) having ≤ 95% allele call rate. Similarly, the mean call rate was ≥ 99.1 % for all validation datasets, highlighting the high typeability of *K. pneumoniae* genomes when using the Kpn-cgMLST scheme.

**Table 1.**
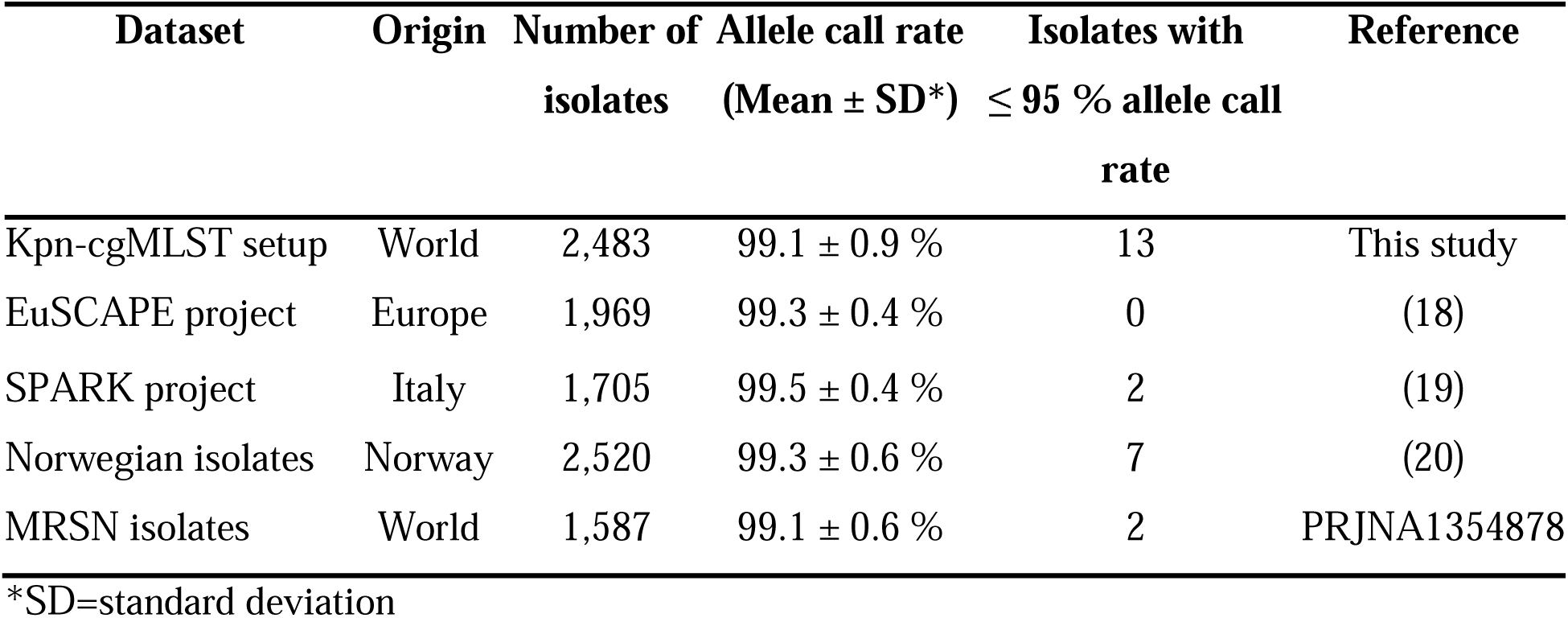
The allele call rate of the isolates of the tested datasets for the Kpn-cgMLST scheme.

| Dataset | Origin | Number of isolates | Allele call rate (Mean ± SD*) | Isolates with $\leq 95$ % allele call rate | Reference |
| --- | --- | --- | --- | --- | --- |
| Kpn-cgMLST setup | World | 2,483 | 99.1 ± 0.9 % | 13 | This study |
| EuSCAPE project | Europe | 1,969 | 99.3 ± 0.4 % | 0 | (18) |
| SPARK project | Italy | 1,705 | 99.5 ± 0.4 % | 2 | (19) |
| Norwegian isolates | Norway | 2,520 | 99.3 ± 0.6 % | 7 | (20) |
| MRSN isolates | World | 1,587 | 99.1 ± 0.6 % | 2 | PRJNA1354878 |

\*SD=standard deviation

The pairwise allelic distances between the isolates were compared when genotyping (i) with only the KpSC-cgMLST (n=629), the MLST (n=7), and the rMLST (n=53) loci or (ii) with the addition of the Kpn-cgMLST loci (3434 in total; in this case, the 7 MLST loci were excluded to avoid redundancy, as the full coding sequences corresponding to these loci are part of the Kpn-cgMLST scheme). The mean pairwise allelic distances between the isolates without and with the Kpn-cgMLST loci were equal to 509 ± 84 and 2856 ± 454, respectively (**Fig. 2A**), constituting an average of 5.6-fold increase. The effect of the inclusion of the Kpn-cgMLST loci on allelic distances for closely related isolates (pairwise distances with KpSC, MLST and rMLST ≤ 40) is illustrated on **Fig. 2B** showing that genomes distinguished by <10 KpSC loci can now be differentiated at up to >100 loci, representing a considerable gain of genotyping resolution.

**Fig. 2.**
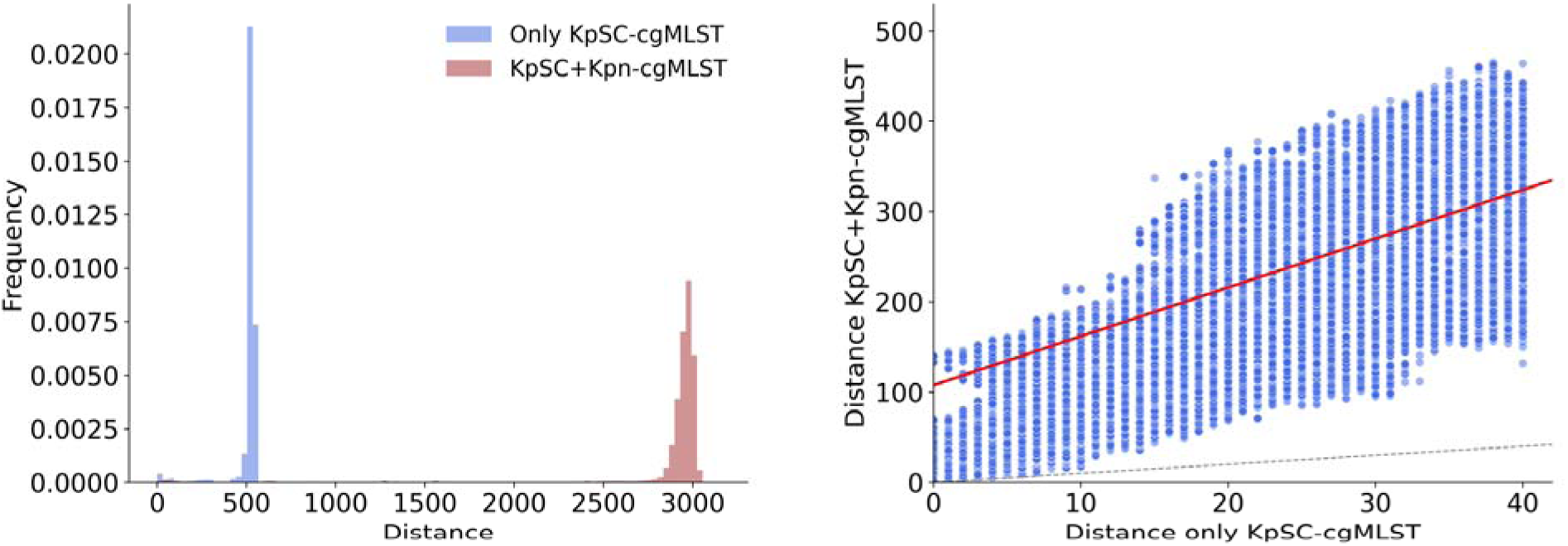
The effect of the inclusion of the Kpn-cgMLST scheme loci on the pairwise allelic distances between the isolates of all the validation datasets. The distances were calculated based on the KpSC-cgMLST (n=629), the MLST (n=7), and the rMLST (n=53) loci (marked as only KpSC-cgMLST on the figure for simplicity) or with the addition of the Kpn-cgMLST loci (n=3434 in total; in this case the 7 MLST loci were excluded). (**A**) The distribution of pairwise distances between the isolates with (red) and without (blue) the new Kpn-cgMLST loci. (**B**) Relationships between the pairwise distances derived with or without the inclusion of the Kpn-cgMLST loci. Allelic distances ≤40 without the Kpn-cgMLST scheme are presented. The dashed line shows the y=x line. The red line shows the linear regression line by applying the least squares method.

### Definition and evaluation of the SL147 cgMLST scheme

We next sought to apply the same strategy at a finer scale of diversity, focusing on the clinically significant SL147 and SL307 *K. pneumoniae* sublineages.

The resulting SL147 cgMLST scheme includes 852 protein-coding loci conserved in *K. pneumoniae* sublineage SL147. A detailed description of the SL147 cgMLST scheme can be found in **Supplementary Material Text 1**. The loci characteristics are presented in **Supplementary Figure S5** and **Supplementary Table S3**. The allele call rates on the setup and validation datasets are shown in **Supplementary Figure S6**.

As expected, the pairwise distances between the validation isolates increased further with the inclusion of the SL147-cgMLST loci, compared to KpSC and Kpn schemes only (**Supplementary Figure S7A**). Most SL147 validation isolates belonged to one the following 4 sequence types: ST147 (n=1112; 79%), ST392 (n=146; 10%), ST4843 (n=46; 3%), ST273 (n=43; 3%), and the occurrence of peaks in the distributions of the pairwise distances could be attributed to the intra- and inter-ST pairs (**Supplementary Figure S7B - D**). Compared to the KpSC+Kpn cgMLST loci, the addition of the sublineage SL147-cgMLST loci increased th pairwise distances between isolates by an average of 20% (**Supplementary Figure S7F**). The SL147-cgMLST loci, together with the Kpn-cgMLST (n=2752), KpSC-cgMLST (n=629) and the rMLST (n=53) loci, sum up to 4286 loci, corresponding to ∼83% of the protein coding sequences of the *K. pneumoniae* NTUH-K2044 reference genome. We refer to the sum of these loci as the cumulative cgMLST scheme for SL147 (hereafter, the cSL147 scheme), which can be used for cgMLST genotyping of *K. pneumoniae* SL147 isolates.

### Genomic investigation of an SL147 outbreak dataset using the cumulative cgMLST strategy

The cumulative cgMLST strategy was used to revisit a previously published *K. pneumoniae* ST147 outbreak dataset (28). By employing the cSL147 scheme, the cumulative cgMLST analysis displayed high performance, as the pairwise allelic distances between the outbreak isolates showed strong concordance with whole genome SNP distances, yielding a Pearson correlation coefficient of 0.97 and a regression slope of 0.84 (**Fig. 3**). With both SNP and cumulative cgMLST analyses, pairwise distances within the previously reported clades A and B of the outbreak were <20. To compare the phylogenetic structures obtained by the two methods, the isolates were clustered using single linkage clustering with a cut-off of 4 based on their SNP distances, which resulted in 21 clusters. These clusters were overlaid onto both the SNP phylogenetic tree (**Fig. 3C**) and the cumulative-cgMLST-based cSL147 minimum spanning tree (**Fig. 3D**). The adjusted rand index (ARI) between the SNP and cumulative cgMLST single linkage clustering labels with a threshold of 4 was 0.95, reflecting the agreement in clustering between these two approaches. The structure of the tree generated using only the rMLST, KpSC and Kpn schemes closely resembled that of the cSL147 scheme, but with a few structural differences and lower pairwise distances. In contrast, as expected, the tree based solely on the KpSC scheme was more divergent and less informative, with many clusters being indistinguishable (**Supplementary Figure S8**). Collectively, these findings indicate that the cumulative cgMLST strategy implementation for the *K. pneumoniae* sublineage SL147 exhibits high discriminatory power and phylogenetic concordance with whole genome SNPs.

**Fig. 3.**
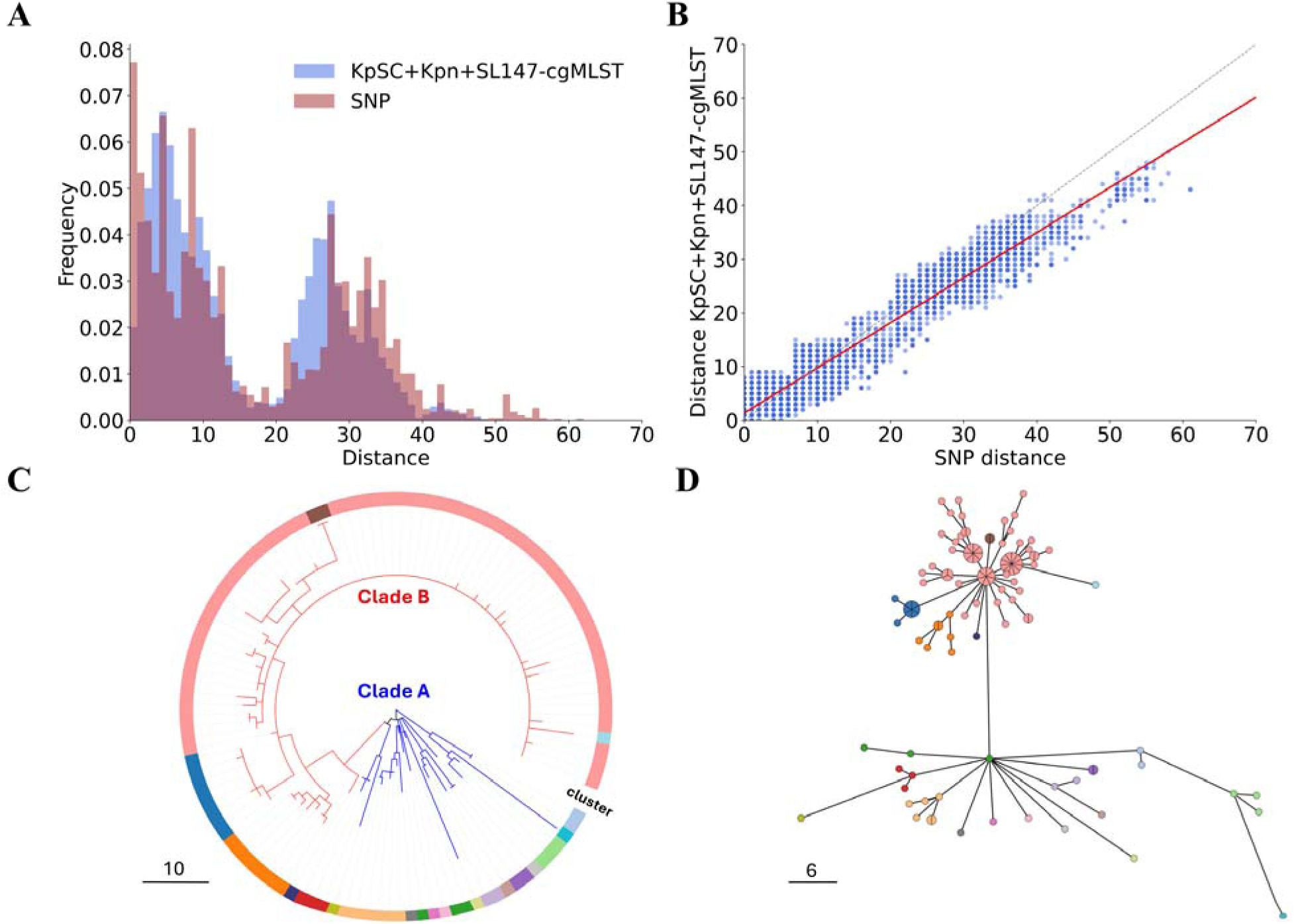
Pairwise cgMLST and SNPs distances between n=114 *K. pneumoniae* SL147 outbreak isolates from dataset 11. Cumulative cgMLST distances were calculated using all loci from the KpSC (n=629 loci), rMLST (n=53 loci), Kpn (n=2,752 loci), and SL147 (n=852 loci) schemes, totaling 4,286 loci. (**A**) Distribution of pairwise distances between isolates based on cumulative cgMLST (blue) and whole-genome SNPs (red). (**B**) Relationships between distances derived from cumulative cgMLST and SNP analyses; the red line indicates the linear regression between the two methods and the dashed line represents y=x. (**C**) SNP-based phylogenetic tree. (**D**) Minimum spanning tree based on cumulative cgMLST distances. The branch colors applied to the SNP-based phylogenetic tree correspond to the clade of the sample (clade A: blue; or clade B: red). The colors applied to both trees correspond to single linkage clusters defined using the whole genome SNP distances, with a cut-off value of 4 SNPs.

### Definition and evaluation of the SL307 cgMLST scheme

Following the same strategy, we implemented an iteration of the cumulative cgMLST strategy for the SL307 sublineage of *K. pneumoniae*. The additional SL307 cgMLST loci comprise 947 protein-coding genes (**Supplementary Material Text 2**). The loci characteristics are presented in **Supplementary Figure S9** and **Supplementary Table S4.** The allele call rates on the setup and validation datasets are shown in **Supplementary Figure S10.**

The cumulative cgMLST scheme (cSL307 scheme) combines the SL307 (n=947), the Kpn (n=2752), the KpSC (n=629), and the rMLST (n=53) loci, totaling 4381 loci and constituting ∼85% of the protein coding sequences of the *K. pneumoniae* NTUH-K2044 reference genome.

As expected, when compared with the higher level cgMLST schemes, the pairwise allelic distances between the SL307 isolates increased with the inclusion of the SL307-cgMLST loci (**Supplementary Figure S11A**); an average increase of 30% was noted (**Supplementary Figure S11C**).

### Genomic investigation of an SL307 outbreak dataset using the cumulative cgMLST strategy

To test the performance of the cumulative cgMLST strategy for outbreak investigation, we used an ST307 outbreak dataset from a single U.S. hospital, which includes 13 isolates from 9 patients and 35 isolates from contaminated sink drains in the hospital. For the clinical isolates, between June 2019 and January 2025, 1 to 3 new patients were detected every year (**Supplementary Table S5**), and cases were dispersed on 7 wards/clinics on four hospital floors. For the environmental swabs, the majority (91%) were taken on a monthly to quarterly basis at the peak of the outbreak, between December 2022-2023, and reservoirs were persistently identified on two floors and wards from where patients were also detected (**Supplementary Table S5**). The cumulative cgMLST pairwise distances were correlated with SNP distances, yielding a Pearson correlation coefficient of 0.93 and a regression slope of 0.66 (**Fig. 4**). To compare the phylogenetic structures obtained by the two methods, the isolates were clustered by single linkage clustering with a cut-off of 4, based on their SNP distances, resulting in 10 clusters. These clusters were mapped onto the SNP phylogenetic tree (**Fig. 4C**) and onto the cSL307 minimum spanning tree (**Fig. 4D**). The defined clusters were arranged similarly in the trees generated using both SNPs and cumulative-cgMLST. The highest obtained adjusted rand index between the SNP distance-based labeling (threshold 4) and cumulative cgMLST distance-based labeling (threshold 2) was 0.6. This moderately high ARI value is likely due to the high genetic relatedness among the isolates, which introduces stochasticity in the polymorphic genetic markers, rather than discrepancies between the two methods. The environmental and human isolates were very closely related (**Fig. 4C**). In contrast, the tree generated using only the KpSC scheme exhibited greater divergence and provided less resolution, with many clusters being indiscernible (**Supplementary Figure S12**).

**Fig. 4.**
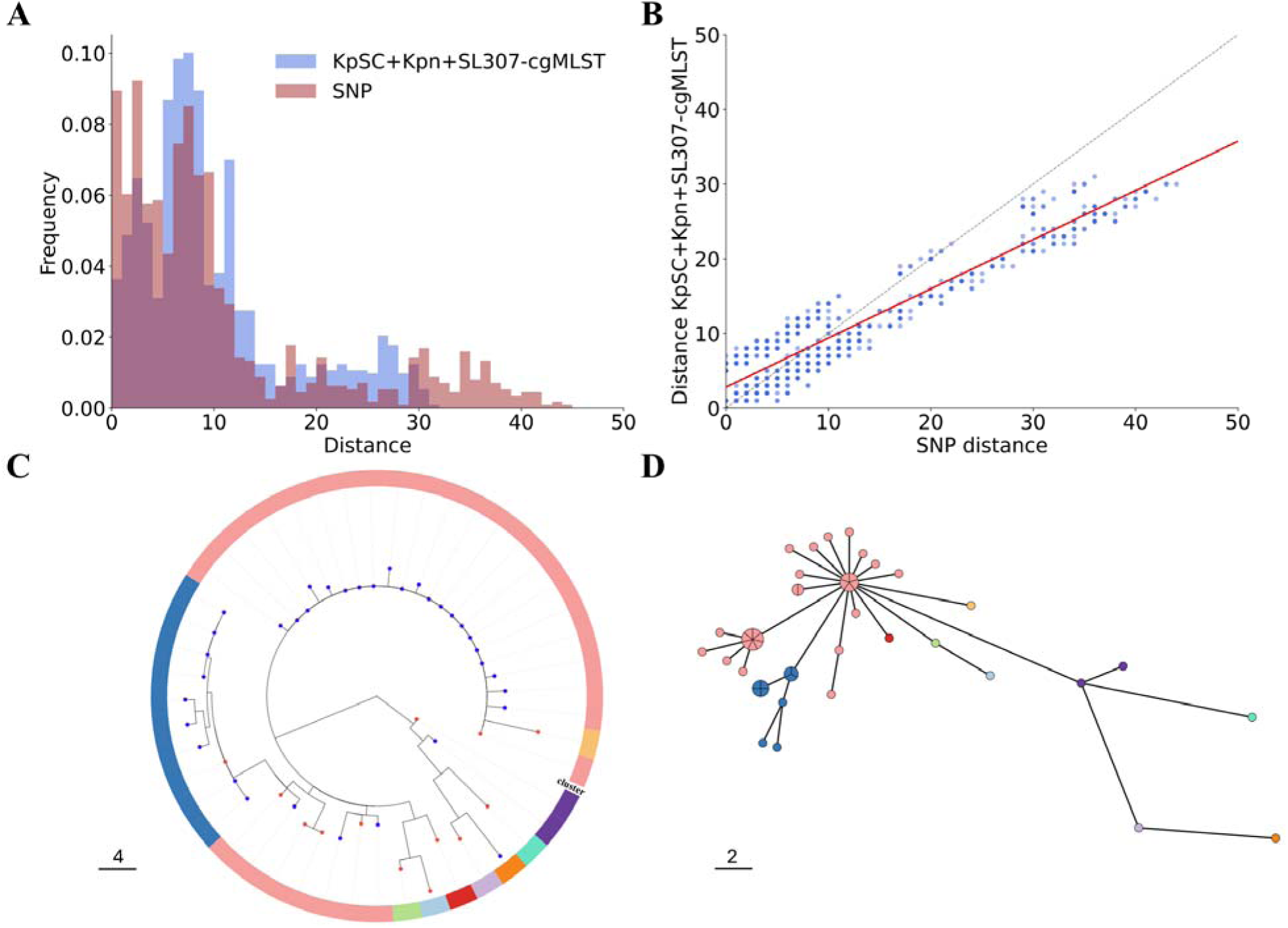
Pairwise cgMLST and SNP distances between n=48 *K. pneumoniae* SL307 outbreak isolates. The cumulative cgMLST distances were calculated based on all loci of the KpSC (n=629), rMLST (n=53), Kpn (n=2752) and the SL307 (n=947) schemes, totaling 4,381 loci. (**A**) Distribution of the pairwise distances between the isolates based on cumulative cgMLST (blue) and whole-genome SNPs (red). (**B**) Relationships between distances derived from cumulative cgMLST and SNP analyses; the red line indicates the linear regression between the two methods and the dashed line represents y=x. (**C**) SNP-based phylogenetic tree. The color of the branch tip circles indicates sample origin (environmental: blue; or human: red). (**D**) Minimum spanning tree based on cumulative cgMLST distances. The colors applied to both trees correspond to single linkage clusters defined using the whole genome SNP distances, with a cut-off value of 4 SNPs.

## Discussion

The use of whole genome sequencing for epidemiological purposes, or genomic epidemiology, has become a central component of infectious disease surveillance and outbreak investigations. The need to combine high resolution bacterial strain genotyping with standardization and reproducibility has so far been difficult to achieve. While whole genome SNP-based approaches can provide high resolution, comparing the resulting genotypes across studies is challenging (2). SNP analysis is typically customized to specific studies/local lineages, which can limit cross-study comparability, although more broadly applicable SNP based genotyping schemes have been developed (40,41). Conversely, the highly standardized cgMLST approach has been limited in its genotyping resolution given its reliance on broadly conserved gene sets that are shared among isolates within species, to be broadly applicable. For example, a typical Enterobacterales genome contains 4,000 to 6,000 genes, but the core genome at the species level is usually restricted to around 2,500 genes. Thus, nearly half of the genomic content is omitted from cgMLST analyses, which results in the overlooking of genetic diversity that could be informative for outbreak source tracing or fine-scale population analysis. Here, we present a strategy to address the resolution gap of cgMLST by cumulating gene loci conserved among increasingly homogeneous phylogenetic groups.

Enhancing cgMLST resolution can be achieved using schemes tailored specifically to the biological entity under study, such as an emerging lineage. These specialized schemes would have limited applicability, as many of their loci would not be present in genomes of other lineages, implying a need to repeat the process for other lineages. Given the high number of such lineages within typical bacterial species, the uncoordinated creation of multiple lineage-specific schemes presents the risk of redundancy of loci across lineage schemes. Further, they would also overlap with those in cgMLST schemes developed for broader phylogenetic levels, leading to redundancy in genotyping analyses and creating challenges in database management.

To address these issues, we propose a structured, systematic process to design new schemes for nested groups. We call this novel approach the cumulative cgMLST strategy. Cumulative cgMLST aims to provide high-resolution genotyping while avoiding redundancy, by the iterative creation of non-overlapping sets of loci conserved within nested phylogenetic entities.

As a proof of concept, we demonstrated the power of the cumulative cgMLST strategy using the *Klebsiella pneumoniae* species complex, for which the widely used KpSC-cgMLST scheme is limited to 629 loci as a result of stringent conservation and synteny inclusion criteria when this scheme was created (7). Specifically, we here developed schemes for *K. pneumoniae sensu stricto* (Kpn-cgMLST), the most prominent member of the KpSC, as well as for its sublineages SL147 and SL307. These schemes comprised all gene loci conserved within the target groups and non-redundant with higher-level schemes. The loci in these sets were integrated within BIGSdb and can be used jointly with the existing *K. pneumoniae* species complex (KpSC) and rMLST schemes (7,14) for high-resolution genotyping (**Supplementary Figure S13**).

The newly created Kpn cgMLST cumulative schemes showed high call rates against geographically and ecologically diverse validation datasets. The combination of the Kpn scheme with the KpSC scheme increased pairwise distances between isolates by an average of 5.6-fold; this scheme can be used for any *K. pneumoniae* sublineage for high resolution genotyping.

As a next iteration, the cumulative cgMLST approach was applied to SL147 and SL307 sublineages. By using outbreak datasets, we showed that the combined schemes lead to nearly matched resolution compared with whole genome SNPs, with the inferred clustering being highly concordant with the phylogenetic relationships identified through SNP analysis. Therefore, cumulative cgMLST appears to represent a highly sensitive and accurate tool for epidemiological analysis and outbreak investigation.

An alternative for high resolution MLST genotyping is the use of pangenome-based schemes and, depending on the gene content of the isolates under analysis, picking a selection of loci from them. Practical implementations of such strategies exist for use at local level (42) but, to our knowledge, their use as central resources enabling cross-study comparability and harmonization remains to be developed. This approach might be attractive, as the gene clusters are initially defined as being distinct. However, the pangenome approach will also require a check for redundancy when novel emerging lineages are considered, as their genomes will comprise novel gene families which need to be distinguished from existing pangenome members. A strategy that uses consecutive MLST schemes of increasing sizes, called multilevel genome typing (MGT), has also been proposed (43). At the level with the highest resolution, MGT employs pangenome-based genotyping. The main aim of the MGT approach is the epidemiological tracing of clones of various longevity, achieved by capturing the genetic identity of bacterial isolates at various resolution levels.

The cumulative cgMLST approach has limitations. Its main bottleneck is the scheme development stage. Although the implementation of automated pipelines, such as chewBBACA (18), can greatly streamline and expedite this process, advanced automation workflows for scheme development are needed so that the cumulative cgMLST approach could be applied to the wide range of existing and emerging bacterial sublineages.

## Conclusions

The cumulative cgMLST strategy demonstrates high discriminatory power across diverse validation outbreak datasets, showing that this approach will enable accurate high-resolution genotyping and effective surveillance and outbreak investigations. The high reproducibility and portability of cumulative cgMLST promises to facilitate the establishment of stable and standardized genotyping systems with high resolution, providing a practical solution to the discrimination/applicability tradeoff of cgMLST. This innovation will therefore enable combining the high discrimination needed for genomic epidemiology and outbreak investigation, with the standardization of genotyping data needed for cross-study comparisons and shared strain nomenclatures (14,43) at the finest subtyping scales.

## List of abbreviations

MLST: multi-locus sequence typing
cgMLST: core genome multi-locus sequence typing
KpSC: *Klebsiella pneumoniae* species complex
SL: sublineage
BIGSdb: Bacterial Isolate Genome Sequence Database
Kpn: *Klebsiella pneumoniae*
SNP: single nucleotide polymorphism
ESBL: extended spectrum beta lactamase
MDR: multidrug-resistant
CG: clonal group
wgMLST: whole genome multi-locus sequence typing
NIPH: non-informative paralogous hits
rMLST: ribosomal MLST
LIN: Life Identification Number

## Declarations

### Availability of data and materials

All the scripts used for this study can be found in the public repository of Institut Pasteur at https://gitlab.pasteur.fr/aovsepia/cumulative_cgMLST.

All genomes, allelic profiles and datasets analyzed during the current study are available at the *Klebsiella pneumoniae* species complex (KpSC) BIGSdb-Pasteur database (https://bigsdb.pasteur.fr/klebsiella/), including:

Dataset 1: A dataset with 2,483 assemblies for the development of the core genome MLST scheme for *K. pneumoniae sensu stricto* (Kpn-cgMLST). A publicly available BIGSdb-Pasteur project (i.e., a browsable list of isolates) entitled “Kpn_cgMLST setup isolates” was created to facilitate retrieval of the data (https://bigsdb.pasteur.fr/cgi-bin/bigsdb/bigsdb.pl?db=pubmlst_klebsiella_isolates&page=query&project_list=184&submit=1).

Dataset 2. A dataset with 1,069 assemblies for the development of the cgMLST scheme for *K. pneumoniae* sublineage SL147. A publicly available BIGSdb-Pasteur project entitled “SL147_cgMLST setup isolates” was created to facilitate retrieval of the data (https://bigsdb.pasteur.fr/cgi-bin/bigsdb/bigsdb.pl?db=pubmlst_klebsiella_isolates&page=query&project_list=188&submit=1).

Dataset 3. A dataset with 1,080 assemblies for the development of the cgMLST scheme for *K. pneumoniae* sublineage SL307. A publicly available BIGSdb-Pasteur project entitled “SL307_cgMLST setup isolates” was created to facilitate retrieval of the data (https://bigsdb.pasteur.fr/cgi-bin/bigsdb/bigsdb.pl?db=pubmlst_klebsiella_isolates&page=query&project_list=189&submit=1).

### Open access

This research was funded, in whole or in part, by Institut Pasteur, by European Union’s Horizon 2020 research and innovation programme, and by the Gates Foundation. Under the grant conditions of the funders, a Creative Commons Attribution 4.0 Generic License has already been assigned to the Author Accepted Manuscript version that might arise from this submission.

### Author information

#### Contributions

S.B. conceived and designed the research and supervised the study. A.O. contributed to the design and conducted the analyses. F.L. and M.M. conducted the SNP analyses. J.D.B. and M.R.P. provided support to the analyses and to maintenance of the BIGSdb platform. A.O. and S.B. wrote the manuscript. J.D.B., F.L. and M.M. contributed to the revision of the manuscript. All authors read and approved the final manuscript.

## Acknowledgements

We thank Audrey Combary, Brice Raffestin and Bryan Brancotte for BIGSdb-Pasteur maintenance and installing updates, Mario Ramirez (Lisbon University, Portugal) for support and feedback on the project, and Keith Jolley (Oxford University, United Kingdom) for continuous developments of the BIGSdb platform.

## References

1. Riley LW. Laboratory Methods in Molecular Epidemiology: Bacterial Infections. Microbiol Spectr. 2018;6(6):1–23.

2. Schürch AC, Arredondo-Alonso S, Willems RJL, Goering R V. Whole genome sequencing options for bacterial strain typing and epidemiologic analysis based on single nucleotide polymorphism versus gene-by-gene–based approaches. Clinical Microbiology and Infection. 2018;24(4):350–4.

3. Tümmler B. Molecular epidemiology in current times. Environ Microbiol. 2020;22(12):4909–18.

4. Maiden MCJ, Bygraves JA, Feil E, Morelli G, Russell JE, Urwin R, et al. Multilocus sequence typing: A portable approach to the identification of clones within populations of pathogenic microorganisms. Proc Natl Acad Sci USA. 1998;95(6):3140–5.

5. Jolley KA, Maiden MCJ. BIGSdb: Scalable analysis of bacterial genome variation at the population level. BMC Bioinformatics. 2010;11(595):1–11.

6. Maiden MCJ, Rensburg MJJ van, Bray JE, Sarah G. Earle SAF, Jolley KA, McCarthy ND. MLST revisited: the gene-by-gene approach to bacterial. Ecol Lett. 2013;11(10):728–36.

7. Bialek-Davenet S, Criscuolo A, Ailloud F, Passet V, Jones L, Delannoy-Vieillard AS, et al. Genomic definition of hypervirulent and multidrug-resistant Klebsiella pneumoniae clonal groups. Emerg Infect Dis. 2014;20(11):1812–20.

8. Harrison OB, Cehovin A, Skett J, Jolley KA, Massari P, Genco CA, et al. *Neisseria gonorrhoeae* Population Genomics: Use of the Gonococcal Core Genome to Improve Surveillance of Antimicrobial Resistance. Journal of Infectious Diseases. 2020;222(11):1816–25.

9. Suetens C, Latour K, Kärki T, Ricchizzi E, Kinross P, Moro ML, et al. Prevalence of healthcare-associated infections, estimated incidence and composite antimicrobial resistance index in acute care hospitals and long-term care facilities: Results from two european point prevalence surveys, 2016 to 2017. Eurosurveillance. 2018;23(46):1–17.

10. WHO. WHO bacterial priority pathogens list, 2024. Bacterial pathogens of public health importance to guide research, development and strategies to prevent and control antimicrobial resistance. 2024.

11. Taylor AW, Blau DM, Bassat Q, Onyango D, Kotloff KL, Arifeen S El, et al. Initial findings from a novel population-based child mortality surveillance approach: a descriptive study. Lancet Glob Health. 2020;8(7):e909–19.

12. Navon-Venezia S, Kondratyeva K, Carattoli A. *Klebsiella pneumoniae*: A major worldwide source and shuttle for antibiotic resistance. FEMS Microbiol Rev. 2017;41(3):252–75.

13. Wyres KL, Lam MMC, Holt KE. Population genomics of *Klebsiella pneumoniae*. Nat Rev Microbiol. 2020;18(6):344–59.

14. Hennart M, Guglielmini J, Bridel S, Maiden MCJ, Jolley KA, Criscuolo A, et al. A Dual Barcoding Approach to Bacterial Strain Nomenclature: Genomic Taxonomy of Klebsiella pneumoniae Strains. Mol Biol Evol. 2022;39(7).

15. Wyres KL, Wick RR, Judd LM, Froumine R, Tokolyi A, Gorrie CL, et al. Distinct evolutionary dynamics of horizontal gene transfer in drug resistant and virulent clones of *Klebsiella pneumoniae*. PLoS Genet. 2019;15(4):1–25.

16. Fostervold A, Hetland MAK, Bakksjø R, Bernhoff E, Holt KE, Samuelsen Ø, et al. A nationwide genomic study of clinical *Klebsiella pneumoniae* in Norway 2001-15: Introduction and spread of ESBLs facilitated by clonal groups CG15 and CG307. Journal of Antimicrobial Chemotherapy. 2022 Mar 1;77(3):665–74.

17. Long SW, Olsen RJ, Eagar TN, Beres SB, Zhao P, Davis JJ, et al. Population genomic analysis of 1,777 extended-spectrum beta-lactamase-producing *Klebsiella pneumoniae* isolates, Houston, Texas: Unexpected abundance of clonal group 307. mBio. 2017;8(3).

18. Silva M, Machado MP, Silva DN, Rossi M, Moran-Gilad J, Santos S, et al. chewBBACA: A complete suite for gene-by-gene schema creation and strain identification. Microb Genom. 2018;4(3):1–7.

19. Kennedy RJ. Prodigal training files for K. pneumoniae used in chewBBACA. https://github.com/B-UMMI/chewBBACA/blob/master/CHEWBBACA/prodigal_training_files/Klebsiella_pneumoniae.trn

20. Jolley KA, Bliss CM, Bennett JS, Bratcher HB, Brehony C, Colles FM, et al. Ribosomal multilocus sequence typing: Universal characterization of bacteria from domain to strain. Microbiology (N Y). 2012;158(4):1005–15.

21. Criscuolo A. FASTQ files to de novo assembly. https://gitlab.pasteur.fr/GIPhy/fq2dna

22. Schwengers O, Jelonek L, Dieckmann MA, Beyvers S, Blom J, Goesmann A. Bakta: Rapid and standardized annotation of bacterial genomes via alignment-free sequence identification. Microb Genom. 2021;7(11).

23. Gurevich A, Saveliev V, Vyahhi N, Tesler G. QUAST: Quality assessment tool for genome assemblies. Bioinformatics. 2013;29(8):1072–5.

24. KlebNET. Whole genome sequencing data requirements for Klebsiella species complex. https://bigsdb.pasteur.fr/klebsiella/genomes-quality-criteria/

25. Grundmann H, Glasner C, Albiger B, Aanensen DM, Tomlinson CT, Andrasević AT, et al. Occurrence of carbapenemase-producing *Klebsiella pneumoniae* and *Escherichia coli* in the European survey of carbapenemase-producing *Enterobacteriaceae* (EuSCAPE): a prospective, multinational study. Lancet Infect Dis. 2017;17(2):153–63.

26. Thorpe HA, Booton R, Kallonen T, Gibbon MJ, Couto N, Passet V, et al. A large-scale genomic snapshot of Klebsiella spp. isolates in Northern Italy reveals limited transmission between clinical and non-clinical settings. Nat Microbiol. 2022;7(12):2054–67.

27. Hetland MAK, Winkler MA, Kaspersen HP, Håkonsholm F, Bakksjø RJ, Bernhoff E, et al. A genome-wide One Health study of Klebsiella pneumoniae in Norway reveals overlapping populations but few recent transmission events across reservoirs. Genome Med. 2025;17(1).

28. Martin MJ, Corey BW, Sannio F, Hall LR, Macdonald U, Jones BT, et al. Anatomy of an extensively drug-resistant *Klebsiella pneumoniae* outbreak in Tuscany, Italy. Proceedings of the National Academy of Science. 2021;118(48):1–8.

29. Bushnell B. https://github.com/BioInfoTools/BBMap/blob/master/sh/bbduk2.sh

30. Wood DE, Lu J, Langmead B. Improved metagenomic analysis with Kraken 2. Genome Biol. 2019 Nov 28;20(1).

31. Seemann T. De novo genome assemblies using shovill https://github.com/tseemann/shovill

32. Bushnell B. https://github.com/BioInfoTools/BBMap

33. Seemann T. Snippy. https://github.com/tseemann/snippy

34. Walker BJ, Abeel T, Shea T, Priest M, Abouelliel A, Sakthikumar S, et al. Pilon: An integrated tool for comprehensive microbial variant detection and genome assembly improvement. PLoS One. 2014;9(11).

35. Croucher NJ, Page AJ, Connor TR, Delaney AJ, Keane JA, Bentley SD, et al. Rapid phylogenetic analysis of large samples of recombinant bacterial whole genome sequences using Gubbins. Nucleic Acids Res. 2015;43(3):e15.

36. Seemann T. https://github.com/tseemann/snp-dists

37. Kozlov AM, Darriba D, Flouri T, Morel B, Stamatakis A. RAxML-NG: A fast, scalable and user-friendly tool for maximum likelihood phylogenetic inference. Bioinformatics. 2019;35(21):4453–5.

38. Zhou Z, Alikhan NF, Sergeant MJ, Luhmann N, Vaz C, Francisco AP, et al. Grapetree: Visualization of core genomic relationships among 100,000 bacterial pathogens. Genome Res. 2018;28(9):1395–404.

39. Diancourt L, Passet V, Verhoef J, Grimont PAD, Brisse S. Multilocus sequence typing of Klebsiella pneumoniae nosocomial isolates. J Clin Microbiol. 2005;43(8):4178–82.

40. Wong VK, Baker S, Connor TR, Pickard D, Page AJ, Dave J, et al. An extended genotyping framework for *Salmonella enterica* serovar Typhi, the cause of human typhoid. Nat Commun. 2016;7:1–11.

41. Hawkey J, Paranagama K, Baker KS, Bengtsson RJ, Weill FX, Thomson NR, et al. Global population structure and genotyping framework for genomic surveillance of the major dysentery pathogen, *Shigella sonnei*. Nat Commun. 2021;12(1).

42. Mixão V, Pinto M, Sobral D, Di Pasquale A, Gomes JP, Borges V. ReporTree: a surveillance-oriented tool to strengthen the linkage between pathogen genetic clusters and epidemiological data. Genome Med. 2023 Dec 1;15(1).

43. Payne M, Kaur S, Wang Q, Hennessy D, Luo L, Octavia S, et al. Multilevel genome typing: Genomics-guided scalable resolution typing of microbial pathogens. Eurosurveillance. 2020;25(20):1–14.

