## Supplementary Appendix for "Cumulative cgMLST provides increased discrimination of nested phylogenetic groups"

### Cumulative cgMLST schemes for increased discrimination of nested phylogenetic groups

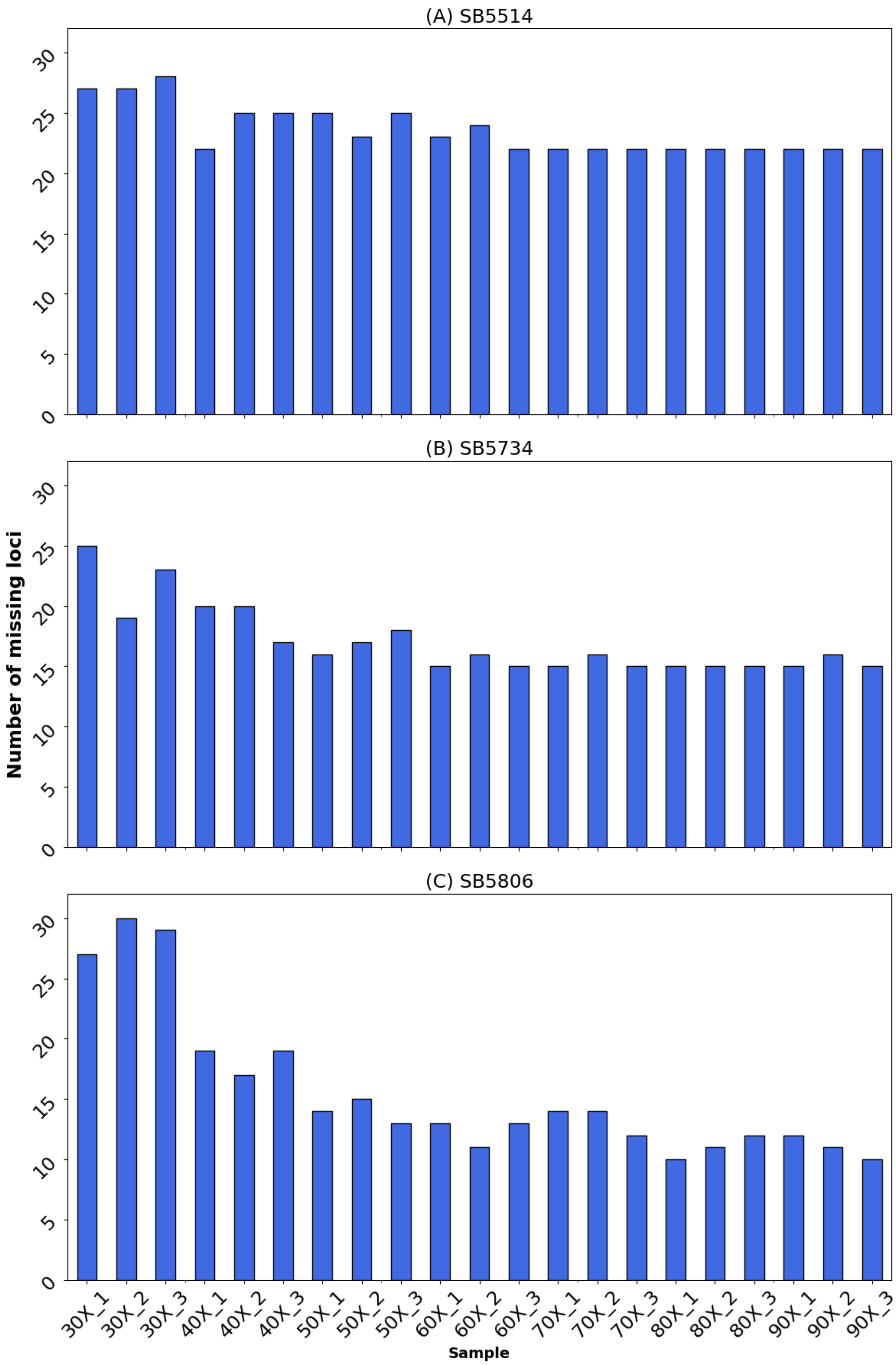

**Figure S1**. Number of missing alleles out of the 2,752 loci of the Kpn-cgMLST scheme for the BIGSdb-Pasteur isolates SB5514 (A), SB5734 (B), SB5806 (C) at 30X to 90X genome coverage, in triplicates.

**Table S1**. The number of isolates collected from each country by the Multidrug-Resistant Organism Repository and Surveillance Network from 2003 until 2021 (dataset 8).

| Country | Number of samples |
|---|---|
| Thailand | 363 |
| Georgia | 359 |
| Peru | 305 |
| Jordan | 218 |
| Kenya | 159 |
| USA | 97 |
| Uganda | 36 |
| Philippines | 25 |
| Ukraine | 7 |
| Germany | 5 |
| Turkey | 4 |
| Italy | 3 |
| Israel | 2 |
| South Korea | 2 |
| Afghanistan | 1 |
| Guam | 1 |

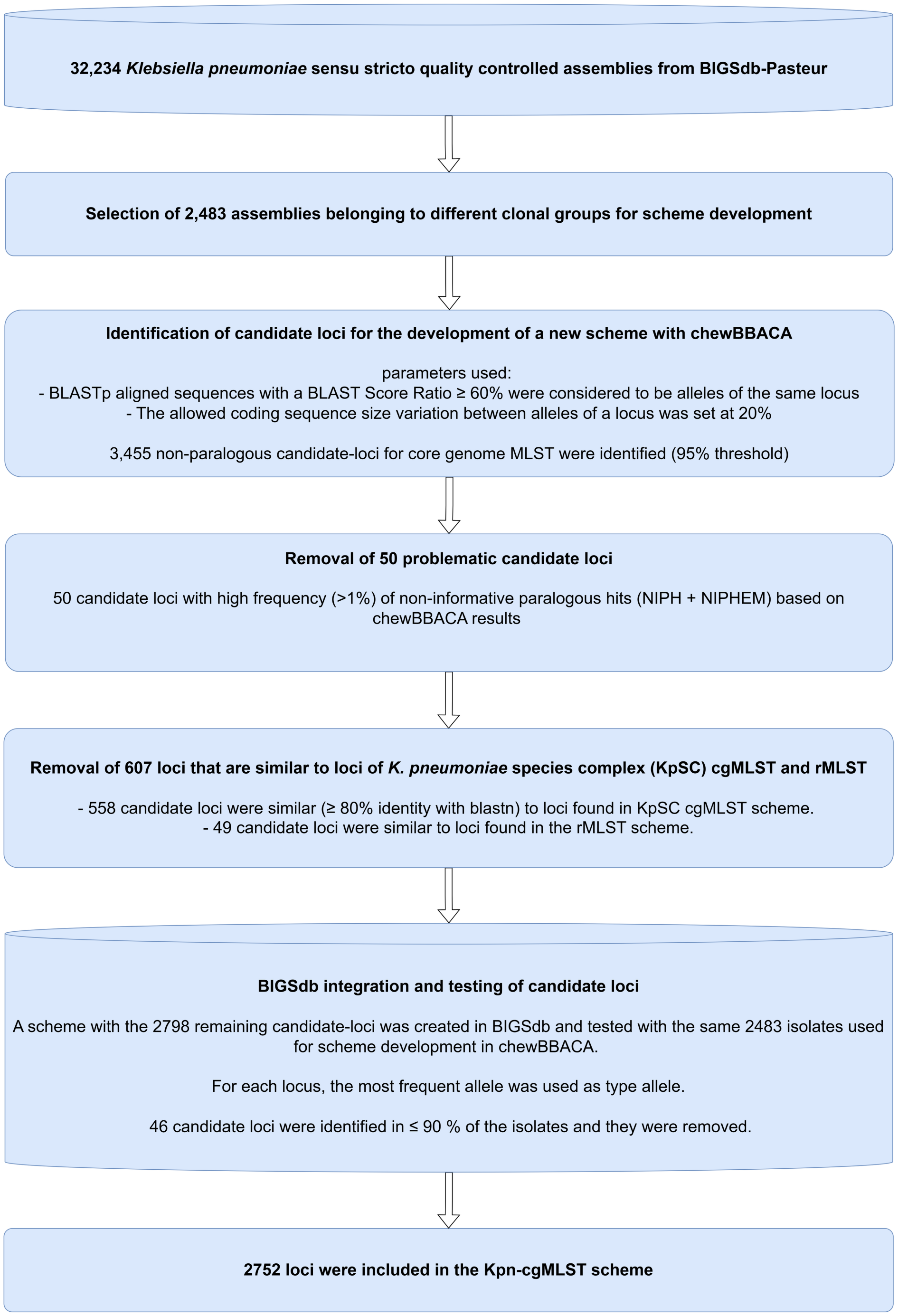

**Figure S2.** Schematic presentation of the main steps followed for the development of the core genome MLST scheme for *Klebsiella pneumoniae sensu stricto* (Kpn-cgMLST).

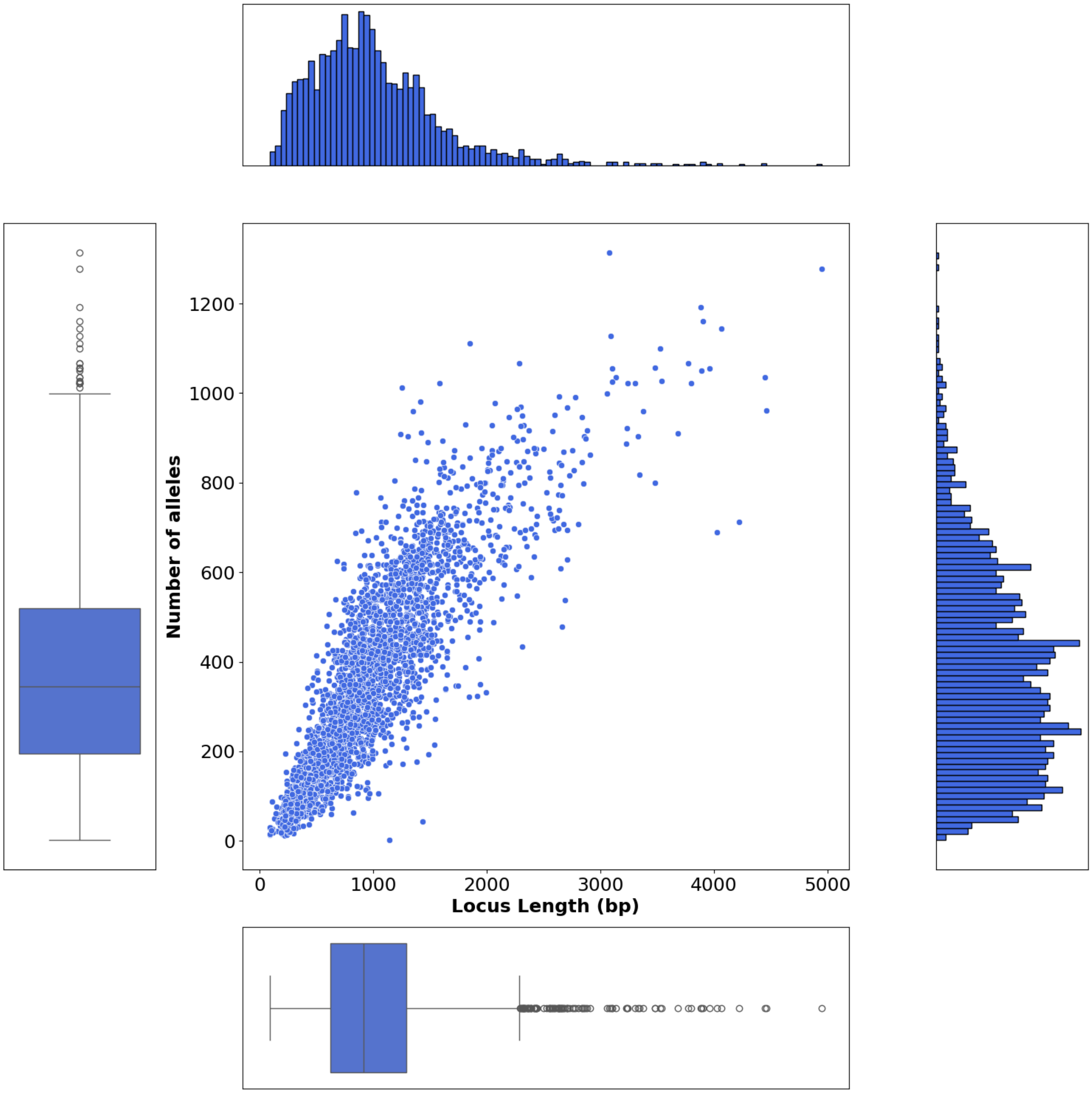

**Figure S3**. 2D-plot showing the relationships between the locus length (bp) and the number of alleles, for all the Kpn-cgMLST loci (n=2752). As locus length, the length of the type allele was used. At the central panel, each point represents a locus; the x axis corresponds to the locus length and the y axis to the number of alleles of that locus. At top/bottom and right/left panels the distributions (100 bins) and the boxplots of each parameter are shown, respectively.

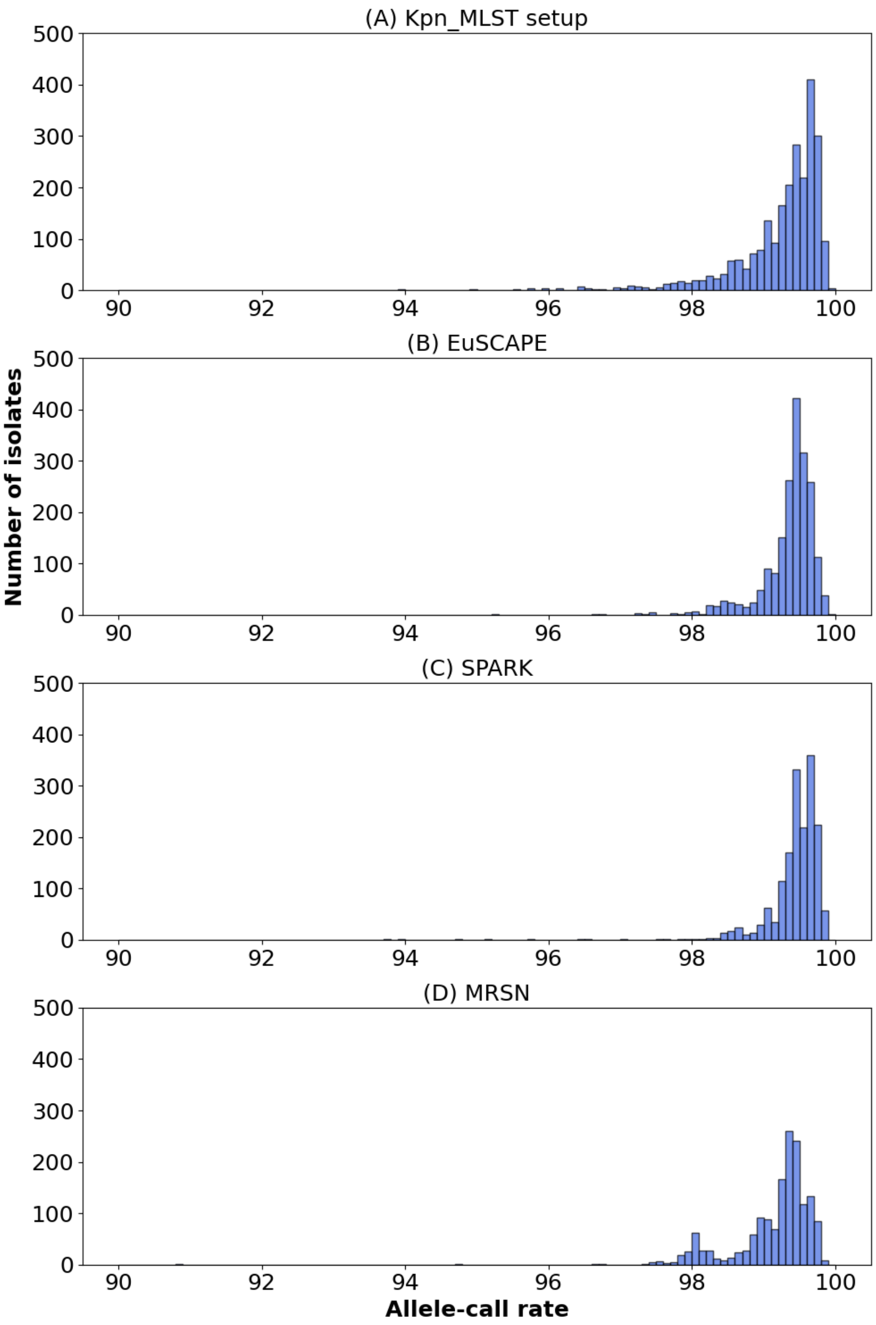

**Figure S4.** The allele call rate (expressed as %) of the Kpn-cgMLST scheme loci (n=2752) in the tested datasets. (A) the 2,483 setup isolates for the creation of the scheme; (B) the 1,969 isolates from the EuSCAPE project; (C) the 1,705 *K. pneumoniae* isolates from the SpARK project; (D) the 1,587 *K. pneumoniae* isolates collected from 2003 until 2021 by the Multidrug-Resistant Organism Repository and Surveillance Network; (E) the 2,520 *K. pneumoniae* isolates collected from multiple ecological sources from 2001 until 2020 in a One Health study in Norway. Bin size=0.1. Note that the X-axis starts at 90%.

**Supplementary Text 1. Description of the SL147 cgMLST scheme**

In the setup dataset of 1,069 genomes, locus lengths ranged from 90 to 4,053 bp, with a total of 1 to 50 alleles per locus, and allele number increasing with locus size (**Supplementary Figure S5**). The locus names, type allele sequences, locus lengths, allowed length ranges, gene names, and gene products are provided in the **Supplementary Table S3**. The mean allele call rate per isolate for the setup isolates was 99.1 ± 1.2 %, with 24 isolates (2.2 %) having ≤ 95% allele call rate (**Supplementary Figure S6A**). The mean allele call rate for the 1,404 isolates of the validation dataset was 99.0 ± 1.6 %, with 42 isolates (3.0 %) having ≤ 95% allele call rate (**Supplementary Figure S6B**).

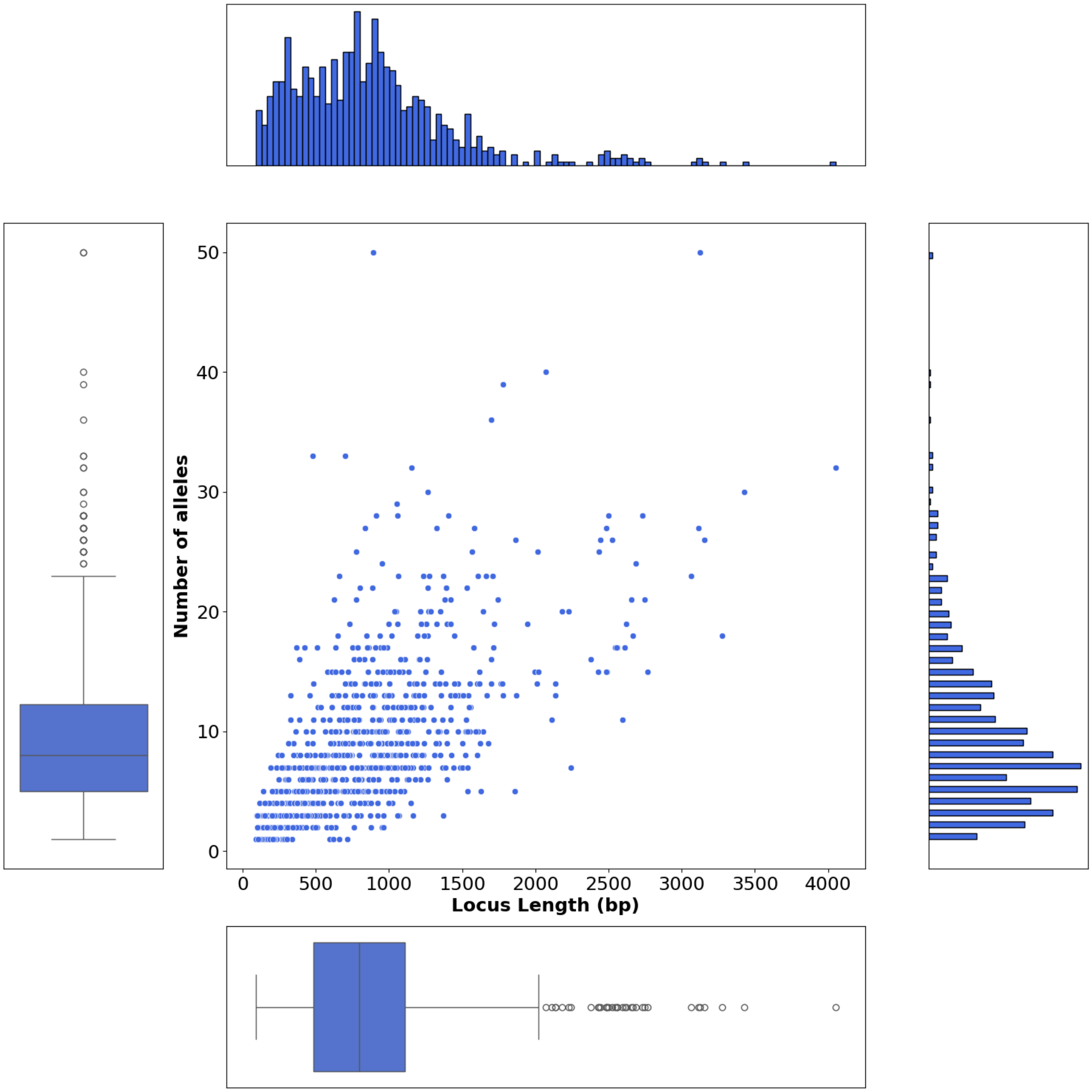

**Figure S5.** 2D-plot showing the relationships between the locus length (bp) and the number of alleles, for all the SL147-cgMLST loci (n=852). As locus length, the length of the type allele was used. At the central panel, each point represents a locus; the x axis corresponds to the locus length and the y axis to the number of alleles of that locus. At top/bottom and right/left panels the distributions (100 bins) and the boxplots of each parameter are shown, respectively.

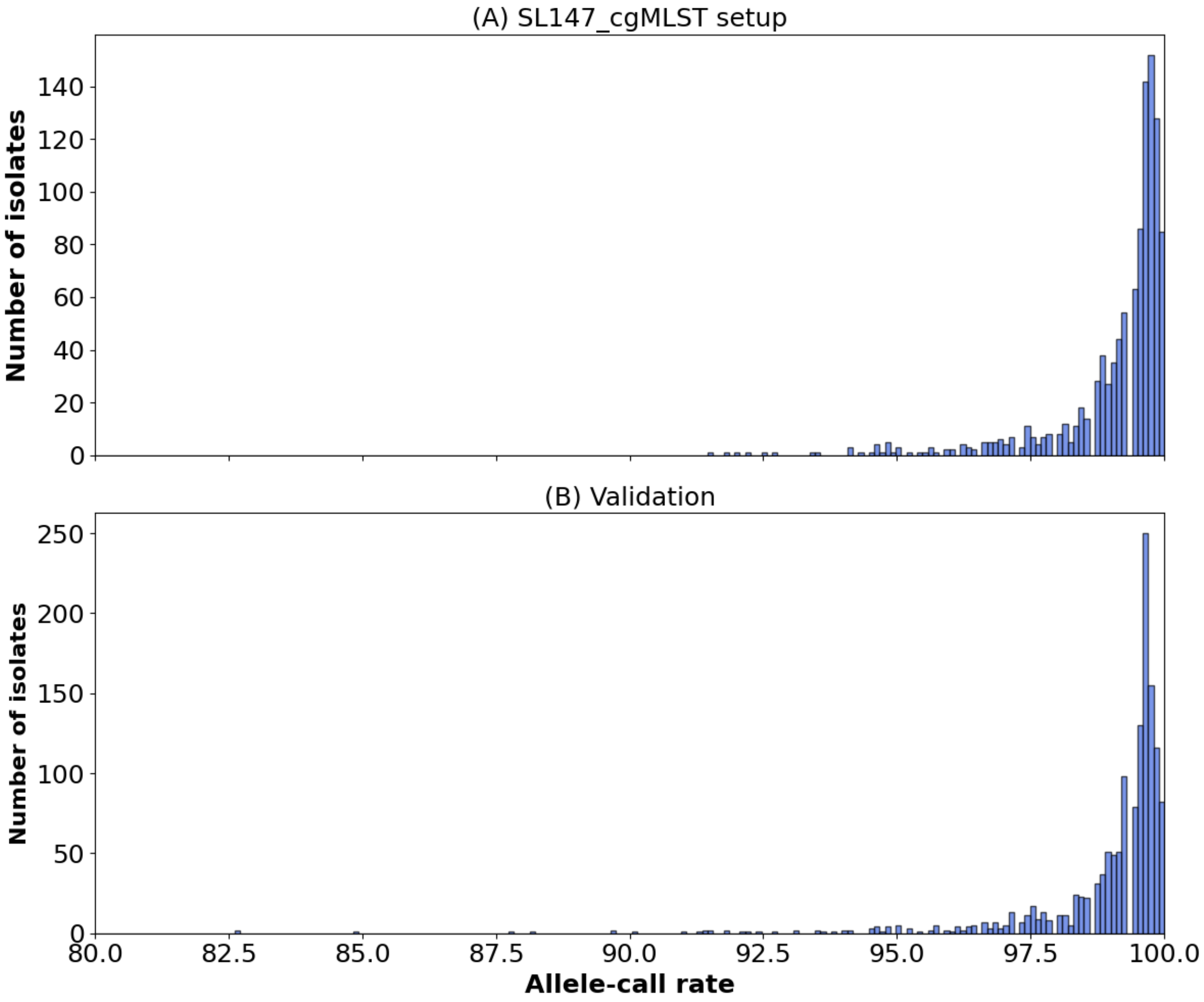

**Figure S6.** The allele call rate (expressed as %) of the SL147 cgMLST scheme loci (n=852) obtained using (A) the 1,069 setup isolates (used for the creation of the SL147 cgMLST scheme); or (B) the 1,404 isolates of the validation dataset. Bin size=0.1. The X-axis starts at 80% for both panels.

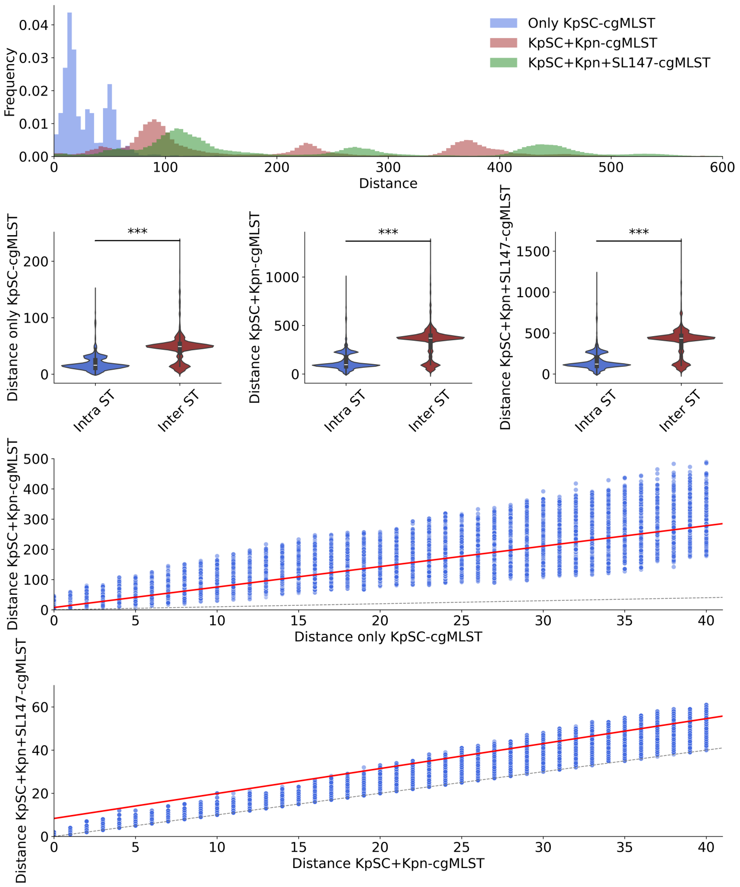

**Figure S7**. The effect of the inclusion of the SL147-specific loci on the pairwise allelic distances between the validation dataset isolates (n=1404). The distances were calculated based on the KpSC (n=629), the MLST (n=7), and the rMLST (n=53) scheme loci together (labeled 'KpSC' on the figure for simplicity), or with the addition of the Kpn (n=2752) and the SL147 (n=852) scheme loci, in which cases the 7 MLST loci were excluded. (**A**) The distribution of pairwise distances between the isolates by using only the KpSC (blue), or additionally the Kpn (red) and then additionally the SL147 (green) cgMLST loci. (**B**, **C**, **D**): Violin plots of the intra- (blue) and inter-ST (red) pairwise distances by using only the KpSC scheme (**B**), or additionally the Kpn scheme (**C**) and then additionally the SL147 scheme (**D**). The nonparametric Mann-Whitney U test was used to determine whether there was a significant difference between the distributions of the intra- and inter- group pairwise allelic distances (*=p≤0.05, **=p≤0.01, and ***=p≤0.001). (**E**) Relationships between pairwise distances derived with or without the inclusion of the Kpn scheme; (**F**) Comparison of pairwise distances with or without the inclusion of the SL147 scheme loci. In E and F, the dashed grey line corresponds to y=x, and the red line corresponds to the linear regression by applying the least squares method.

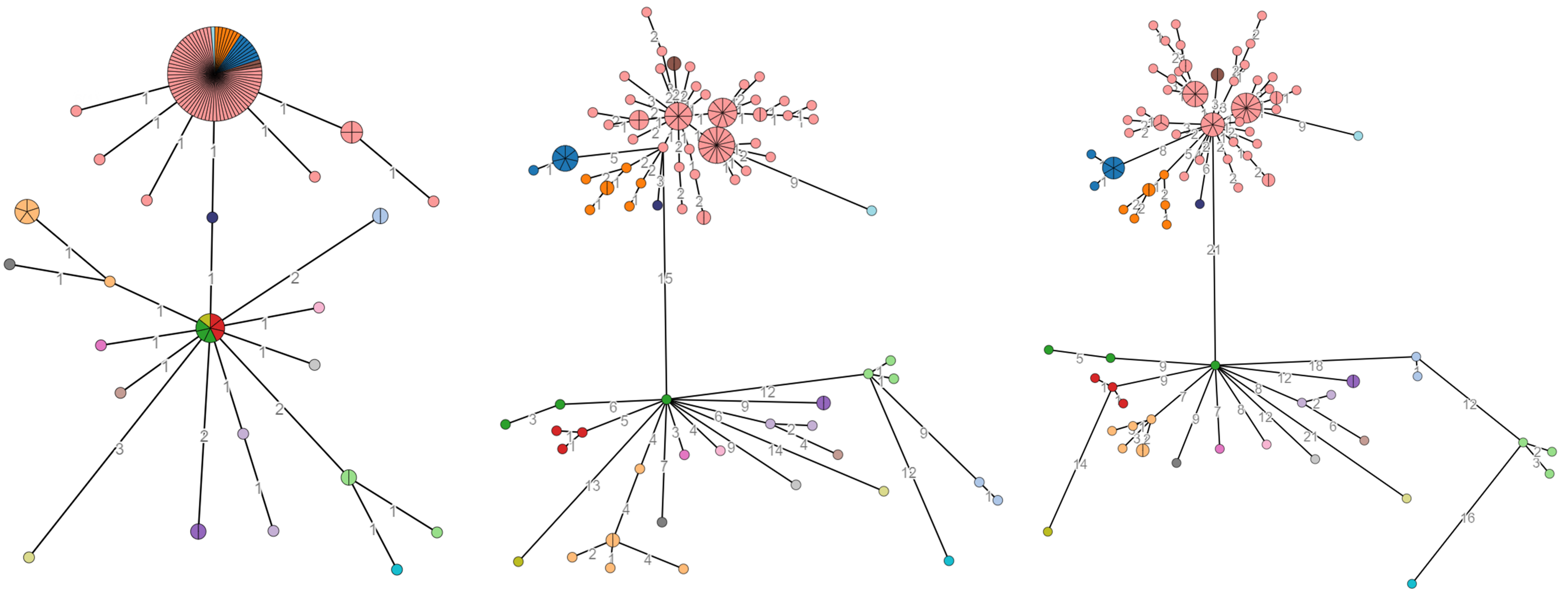

**Figure S8**. Minimum spanning trees based on cgMLST distances between the n=114 SL147 outbreak isolates described in **Methods** (dataset 11). The distances were calculated based on the KpSC scheme (n=629) (plus the MLST (n=7) and the rMLST (n=53) loci) (**A**) or with the addition of the Kpn scheme loci (n=2752) (**B**), or also additionally with the SL147 scheme loci (**C**), in which cases the 7 MLST loci were excluded. The colors applied to the trees correspond to single linkage clusters defined using the whole genome SNP distances, with a cut-off value of 4 SNPs.

**Supplementary text 2. Description of the SL307 cgMLST scheme**

In the reference dataset of 1,080 genomes, locus lengths ranged from 90 to 4,278 bp, with one to 33 alleles per locus, and the number of alleles increased with locus size (**Supplementary Figure S9**). **Supplementary Table S4** provides detailed information on locus names, type allele sequences, locus lengths, allowed length ranges, gene names, and gene products. The mean allele call rate per isolate for the SL307-cgMLST setup isolates was 99.4 ± 1.0 %, with 8 isolates (0.7 %) having ≤ 95% allele call rate (**Supplementary Figure S10A**). The mean allele call rate for the 947 isolates of the validation dataset was 99.3 ± 1.4 %, with 19 isolates (1.9 %) having ≤ 95% allele call rate (**Supplementary Figure S10B**).

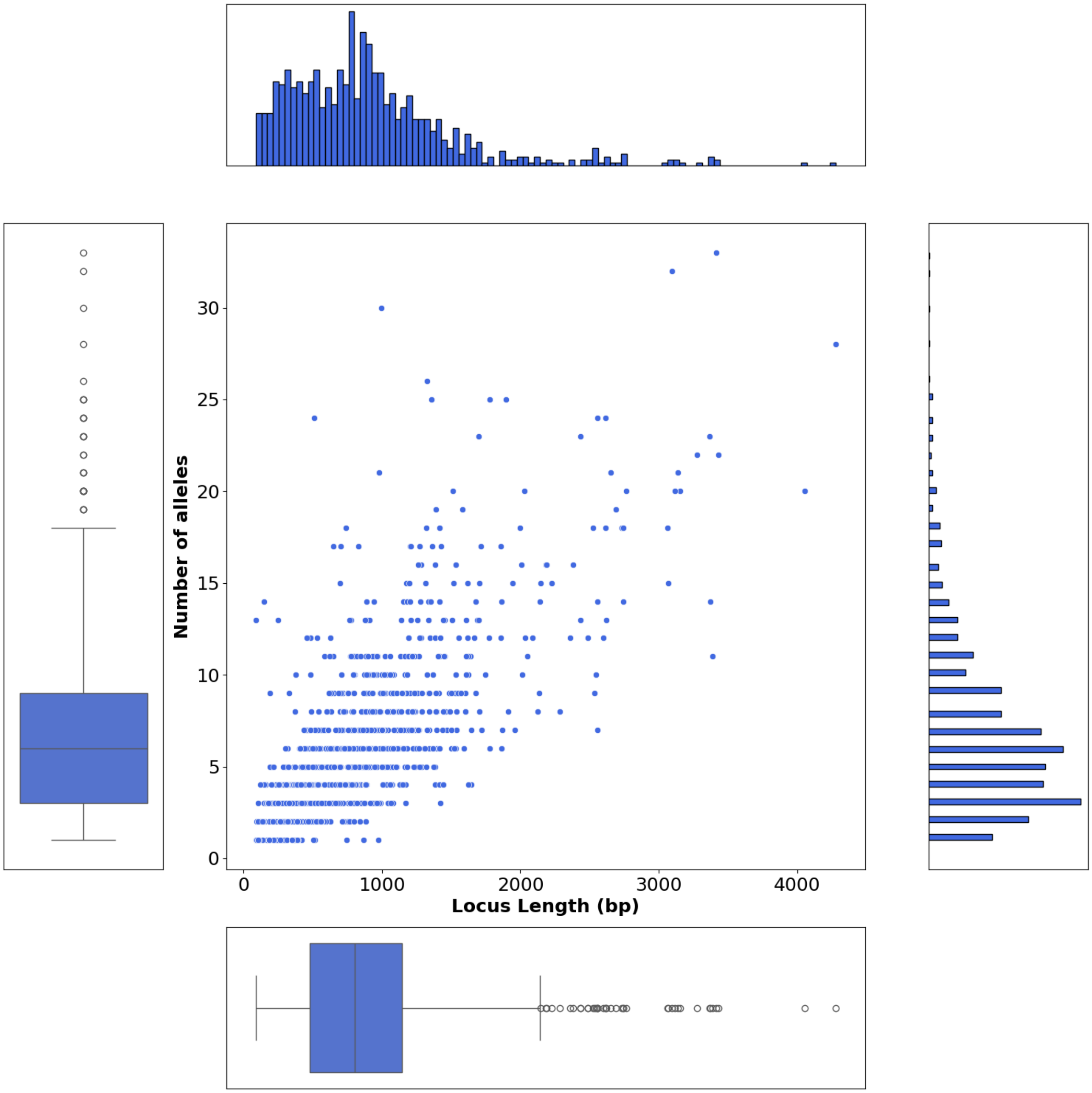

**Figure S9.** 2D-plot showing the relationships between the locus length (bp) and the number of alleles, for all the SL307-cgMLST loci (n=947). As locus length, the length of the type allele was used. At the central panel, each point represents a locus; the x axis corresponds to the locus length and the y axis to the number of alleles of that locus. At top/bottom and right/left panels the distributions (100 bins) and the boxplots of each parameter are shown, respectively.

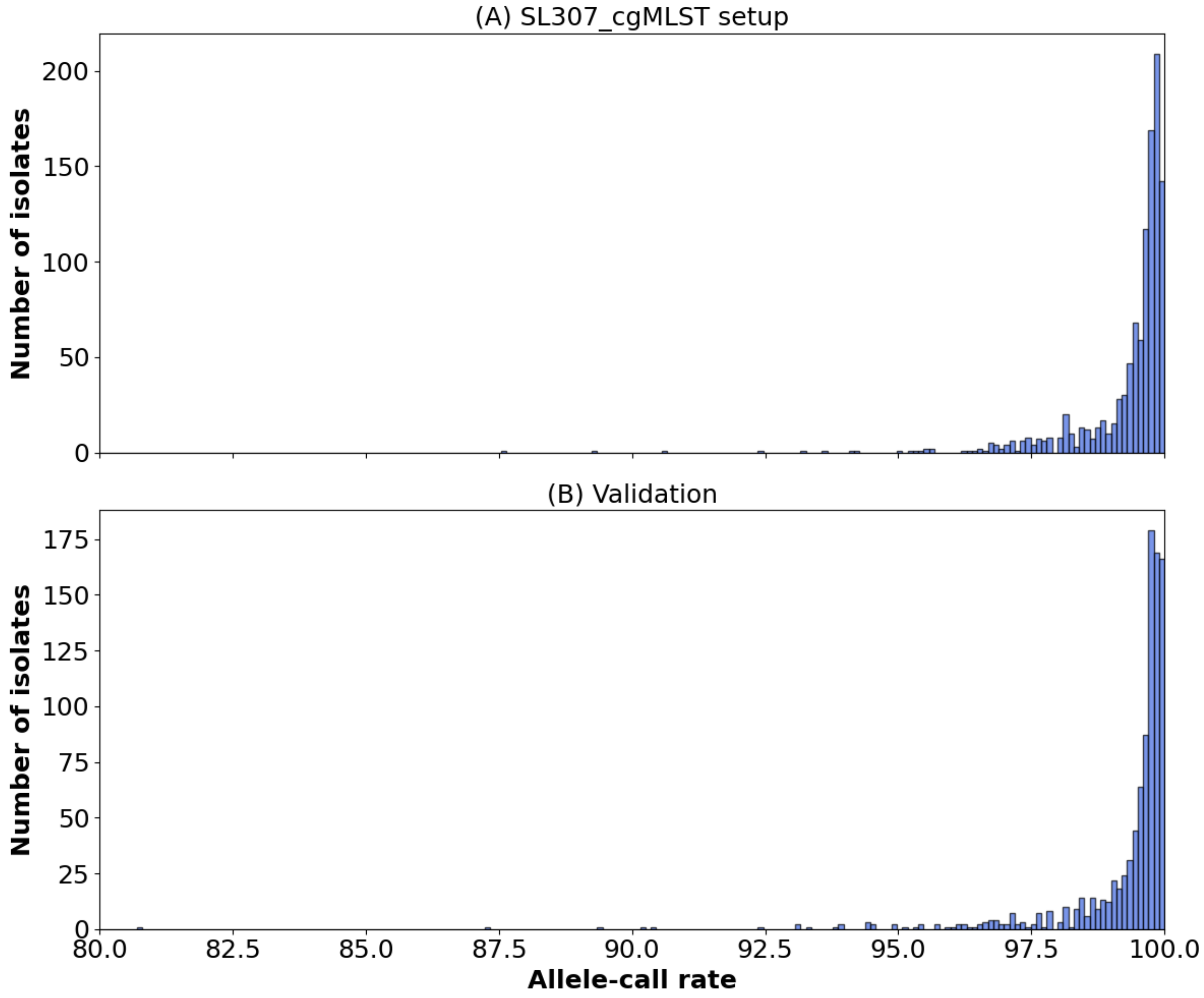

**Figure S10.** The allele-call rate of the SL307-cgMLST scheme (n=947) in the tested dataset. (A) the 1,080 setup isolates for the creation of the SL307 core genome MLST scheme; (B) the 947 isolates of the validation dataset. Bin size=0.1. The X-axis starts at 80% for both panels.

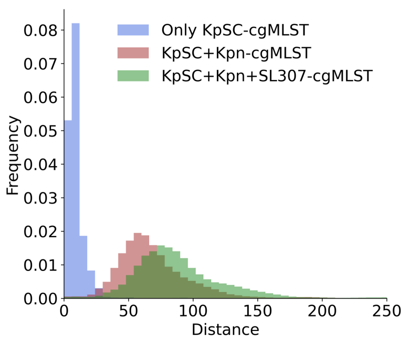

**Figure S11**. The effect of the inclusion of the SL307-specific loci on the pairwise allelic distances between the SL307-cgMLST validation isolates. The distances were calculated based on the KpSC (n=629), the MLST (n=7), and the rMLST (n=53) scheme loci together (labeled 'KpSC' on the figure for simplicity), or with the addition of the Kpn (n=2752) and the SL147 (n=947) scheme loci, in which cases the 7 MLST loci were excluded. (**A**) The distribution of pairwise distances between the isolates by using only the KpSC (blue), or additionally, the Kpn (red) and then additionally the SL307 (green) cgMLST loci. (**B**) Relationships between pairwise distances derived with or without the inclusion of the Kpn scheme; (**C**) Comparison with or without the inclusion of the SL307 scheme loci. In **B** and **C**, the dashed grey line corresponds to y=x, and the red line corresponds to the linear regression by applying the least squares method.

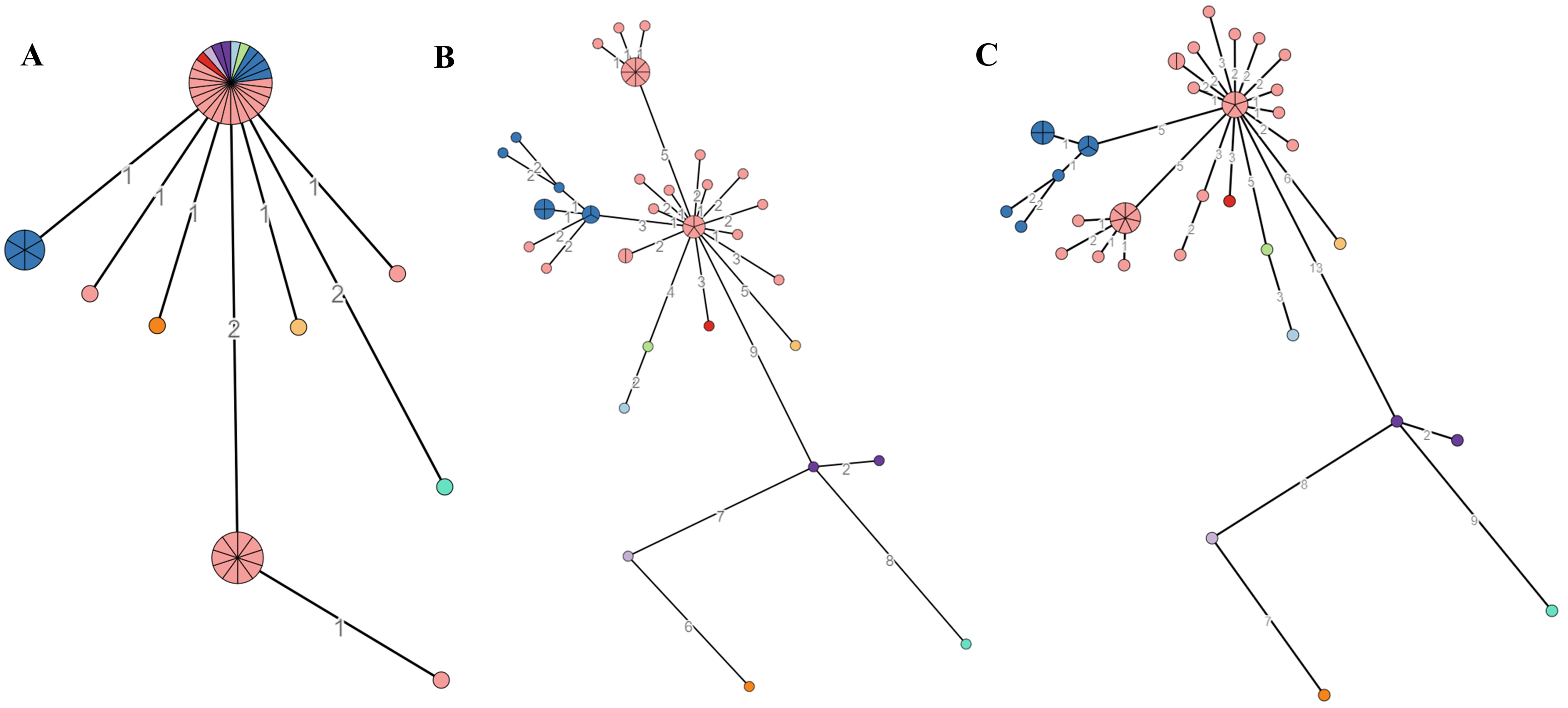

**Figure S12**. Minimum spanning trees based on the cgMLST distances between n=48 *K. pneumoniae* SL307 outbreak isolates described in **Methods** (dataset 12). The distances were calculated based on the KpSC scheme (n=629) (plus the MLST (n=7) and the rMLST (n=53) loci) (**A**) or with the addition of the Kpn scheme loci (n=2752) (**B**), or additionally the SL307 scheme loci (n=947) (**C**), in which cases the 7 MLST loci were excluded. The colors applied to the trees correspond to single linkage clusters defined using the whole genome SNP distances, with a cut-off value of 4 SNPs.

### A. Available schemes and their use for genotyping

| | Species complex schemes | | | Species schemes | Sublineage schemes | |
|---|---|---|---|---|---|---|
| Isolate | MLST | rMLST | KpSC-cgMLST | Kpn-cgMLST | SL147-cgMLST | SL307-cgMLST |
| | | | 629 loci | 2752 loci | 852 loci | 947 loci |
| Any KpSC | ✓ | ✓ | ✓ | | | |
| Any Kpn | | ✓ | ✓ | ✓ | | |
| SL147 | | ✓ | ✓ | ✓ | ✓ | |
| SL307 | | ✓ | ✓ | ✓ | | ✓ |

### B. Selection of schemes in BIGSdb-Pasteur

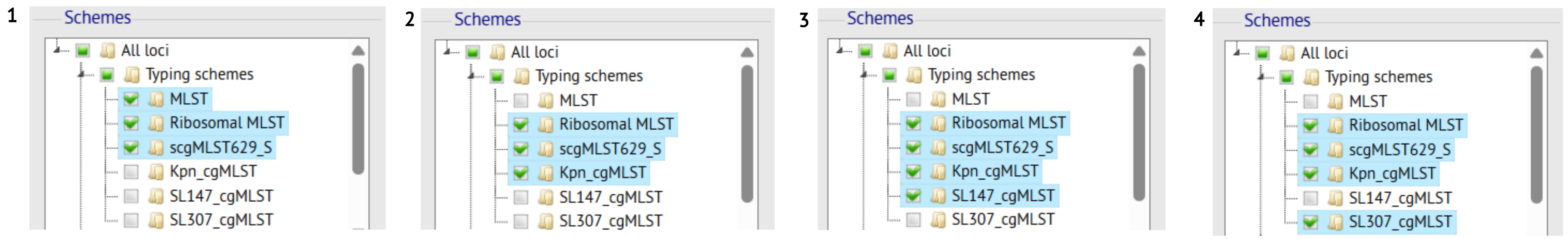

**Figure S13**. (**A**) Available schemes in BIGSdb-Pasteur for the genotyping of *K. pneumoniae* species complex (KpSC) isolates within the framework of cumulative-cgMLST strategy. (**B**) Selection of schemes in the BIGSdb-Pasteur database (https://bigsdb.pasteur.fr/klebsiella/) depending on the isolates under investigation (**1**: any KpSC isolate; **2**: any Kpn isolate; **3**: Kpn SL147 isolates; **4**: Kpn SL307 isolates).
